# Brassinosteroids promote sugar synthesis by inhibiting BIN2 phosphorylation of phosphoenolpyruvate carboxykinase

**DOI:** 10.1101/2025.10.22.683954

**Authors:** Hongliang Zhang, Yalikunjiang Aizezi, Kanako Bessho-Uehara, Ajeet Chaudhary, Cao Son Trinh, Shou-Ling Xu, Zhi-Yong Wang

## Abstract

Sugar is both an essential energy source and the major substrate for cell wall biosynthesis during plant growth, yet how growth-promoting hormones regulate sugar synthesis remains unclear. Here, we show that the brassinosteroids (BRs) promote gluconeogenic and photosynthetic sugar synthesis by activating phosphoenolpyruvate carboxykinase (PCK), which catalyzes the conversion of oxaloacetate to phosphoenolpyruvate, a central step in primary metabolism. Arabidopsis BR-deficient mutants display reduced PCK1 activity and elevated phosphorylation at conserved Ser-62 and Thr-66 residues. BR treatment induces PCK1 dephosphorylation and activation, whereas the GSK3-like kinase BIN2 phosphorylates these sites, altering quaternary structure and inhibiting PCK1. Phospho-blocking mutations of Ser-62/Thr-66 confer BR-independent PCK1 activity and enhance seedling growth, while phosphomimetic mutations reduce PCK1 activity and impair seedling growth and establishment. BR also promotes PCK dephosphorylation and activation in photosynthetic leaves of maize and sorghum. Our study demonstrates that BR regulates primary metabolism via GSK3/BIN2-mediated phosphorylation of PCK, thereby promoting gluconeogenesis and photosynthesis.

## Main

Plant growth consumes sugars for both energy production and cell wall synthesis. Growth-promoting hormones, such as BRs, are known to increase plant growth and biomass accumulation by enhancing the sink strengths. Little is known about whether and how growth-promoting hormones enhance sugar production in source tissues. Primary metabolism channels carbon and nitrogen among carbohydrates, lipids, amino acids, and nucleotides. Regulation of primary metabolic pathways by internal and environmental signals is crucial for maintaining homeostasis and supplying chemical substrates for growth, cell differentiation, and secondary metabolism^1,2^. Understanding how growth hormones regulate primary metabolic pathways is crucial for improving plant productivity and resilience.

BRs are a class of growth-promoting hormones that significantly enhance cell elongation/expansion and biomass accumulation^3–5^. BRs bind to the receptor kinase BRI1 (BRASSINOSTEROID INSENSITIVE 1), initiating a signaling cascade that inactivates BR-INSENSITIVE 2 (BIN2), a glycogen synthase kinase 3 (GSK3)/SHAGGY-like kinase^6–8^. This process leads to the dephosphorylation of BIN2 substrate proteins^9^, including the BRASSINAZOLE RESISTANT 1 (BZR1) family transcription factors^10^. Upon BR inactivation of BIN2, BZR1 is dephosphorylated by PROTEIN PHOSPHATASE 2A (PP2A)^11^, dissociates from the 14-3-3 proteins to move into the nucleus^12^, and activates or represses thousands of target genes by recruiting different chromatin remodelers^13–16^. Nuclear BZR1 interacts with transcription factors of the auxin, gibberellin, and phytochrome pathways to co-regulate growth-promoting gene^17–19^, particularly those involved in cell wall biosynthesis^17,20^. Additionally, BR promotes cell wall synthesis and integrity by releasing BIN2’s phosphorylation/inhibition of cellulose synthase and FERONIA receptor kinase^21,22^.

BR-induced growth consumes sugars for energy and as substrates for cell wall synthesis and is therefore dependent on sugar availability. To maintain cellular homeostasis, BR signaling is modulated by sugar signaling^23^. Sugar signaling via the Target of Rapamycin (TOR) stabilizes BZR1, thereby enabling BR signaling. Under sugar-limiting conditions, inactivation of TOR leads to ubiquitination of BZR1 by the UPL3 ubiquitin ligase followed by degradation by the proteasome and autophagy pathways, leading to growth arrest and improving survival of the stress^24^. Treatment of light-growth seedlings with high levels of sugars also promotes BIN2 phosphorylation of BZR1, thereby inhibiting growth^25^. A recent multi-omics study provided genome-wide evidence for extensive interactions between BR and sugar-TOR signaling pathways^26^. These studies revealed extensive crosstalk between BR and sugar signaling. However, it remains unknown whether BRs promote sugar synthesis to fuel growth.

Proteomic studies have identified BR-regulated phospho-proteins^27,28^, including hundreds of *in vivo* BIN2 substrate proteins^9^. These studies indicate that BRs regulate the phosphorylation of phosphoenolpyruvate carboxykinase 1 (PCK1 or PEPCK1), a key enzyme in primary metabolism including gluconeogenesis and photosynthesis^29,30^. PCK catalyzes the conversion of oxaloacetate (OAA) to phosphoenolpyruvate (PEP) and CO_2_, a rate-limiting step in gluconeogenesis and a key interface between organic acid/amino acid/lipid and sugar metabolism^1,31^. The Arabidopsis genome includes two *PCK* genes, *PCK1* (AT4G37870) and *PCK2* (AT5G65690)^32,33^. While both genes are expressed in various tissues, *PCK1* is expressed at higher levels and in a wider range of tissues^32,34,35^. Arabidopsis PCK1 is essential for converting seed storage lipid into sugar to support post-germination seedling growth before the photosynthetic apparatus is developed. The Arabidopsis *pck1* mutant seedlings exhibit reduced soluble sugar levels, growth impairment, and a low success rate of seedling establishment when grown under extended darkness or weak light conditions^32,36,37^. PCK is also important in vegetative and reproductive tissues^1,34^. Adult plants of Arabidopsis *pck1* mutant show altered sugar and amino acid contents in the vein tissues and display defects in stomata closure^35,38^. In tomato, PCK deficiency affects the germination, growth, and fruit sugar content^39^.

In C4 photosynthesis and Crassulacean acid metabolism (CAM) plants, PCK plays a crucial role in the photosynthetic carbon assimilation, providing CO_2_ for the Calvin-Benson cycle through the decarboxylation of OAA^[40,41]^. PCK also regulates the supply of carbon skeletons for specific metabolic pathways^35^. Recent studies suggest that PCK increases light harvesting plasticity and improves photosynthesis rate under stress conditions^42,43^. In both C3 and C4 plants, PCK activity is increased by light and decreased under shade or dark conditions^31,44,45^. The decreased activity is correlated with increased phosphorylation, suggesting that PCKs are inhibited by phosphorylation^46–48^. However, a causal relationship between phosphorylation and PCK inactivation was not supported by site-directed mutagenesis of the phosphorylation sites. Further, the kinases and phosphatases that regulate the reversible phosphorylation of PCK remain elusive^49^.

Here, we demonstrate that PCK1 is inhibited by the GSK3-like kinase BIN2 and activated through BR-induced dephosphorylation. Extensive *in vitro* and *in vivo* experiments demonstrate that BIN2 phosphorylates PCK1 at two conserved residues, Ser-62 and Thr-66, resulting in quaternary structural changes and inhibition of PCK1 catalytic activity. Mutagenesis of the phosphorylation sites demonstrates their essential function in BR regulation of PCK1 decarboxylase activity and in establishing Arabidopsis seedlings in soil. BR induces PCK1 dephosphorylation and activation not only in Arabidopsis seedlings, where PCK mediates gluconeogenesis, but also in leaves of maize and sorghum, where PCK is inhibited at night but activated during the day to presumably mediate C4 photosynthesis. Our study elucidates a direct link between BR signaling and primary metabolism, enabling BR regulation of sugar production via gluconeogenesis and photosynthesis, thereby ensuring a sufficient sugar supply to meet growth demands in plants. Identifying the mechanistic links between BR signaling and carbon metabolism, particularly C4 photosynthesis, will inform new approaches to improving crop productivity.

## Results

### BRs increase PCK1 activity by inhibiting BIN2

Our proteomic study of BR-regulated phosphoproteins showed that BR induces PCK1 dephosphorylation^27^. Subsequent studies showed dynamic changes in PCK1 phosphorylation following BR treatment^28^, and provided evidence for BIN2 phosphorylation of PCK1^9^. We therefore tested whether BR regulates PCK1 activity. We grew Arabidopsis seedlings in the dark on medium supplemented with varying concentrations of the BR biosynthesis inhibitor propiconazole (PPZ)^50^. PPZ inhibited hypocotyl growth and reduced the PCK1 decarboxylase activity, and the PPZ effects were reversed by brassinolide (the most active form of BR) (Fig. 1a-c; Extended Data Fig. 1a-c). To test whether BR regulates PCK1 through BIN2, we measured PCK1 activity in the dominant *BR-insensitive 2* (*bin2-1*) mutant, in which BIN2 is constitutively active and cannot be inactivated by BR. Like PPZ-treated seedlings, the *bin2-1* seedlings showed short hypocotyls and reduced PCK1 activity (Fig. 1d-f; Extended Data Fig. 1a-c). The phenotypes of *bin2-1* and wild-type seedlings on PPZ were reversed by bikinin, a chemical BIN2 inhibitor^51^ (Extended Data Fig. 1a-c). *PCK1* transcript levels and PCK1 protein levels are not significantly changed by PPZ treatment (Fig. 1g,h). These results indicate that BR signaling increases PCK1 activity via a BIN2-dependent mechanism.

**Fig. 1:**
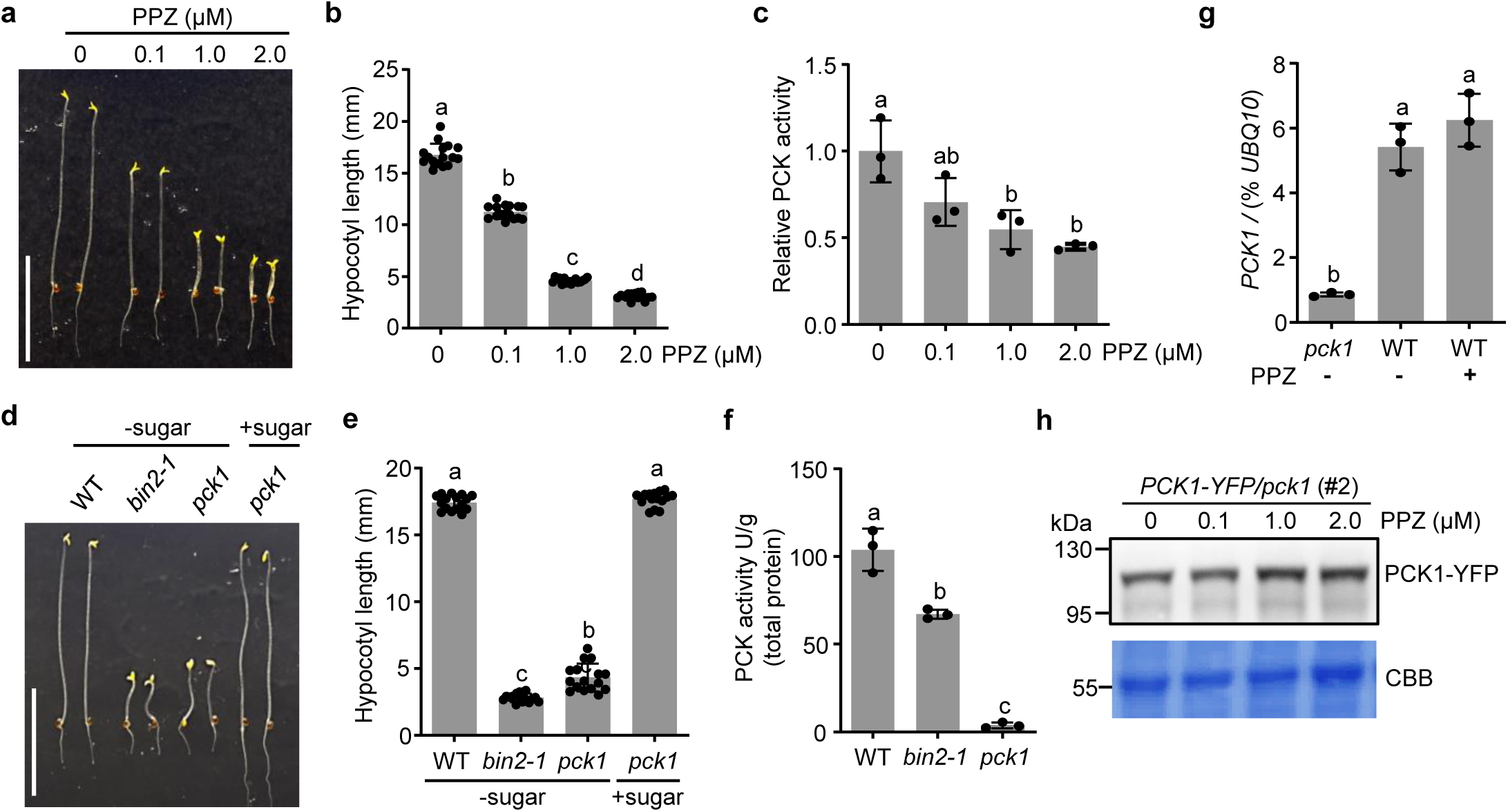
BRs regulate Phosphoenolpyruvate Carboxykinase (PCK) activity through BIN2. **a**-**c,** Wild-type (WT) seedlings were grown on 0.5x MS sugar-free medium containing the indicated concentrations of propiconazole (PPZ) in the dark for 5 days. Representative seedlings were imaged (**a**), hypocotyl length (**b**) and PCK activity (**c**) were measured. **d**-**f,** WT, *bin2-1*, and *pck1* seedlings were grown on 0.5x MS medium with or without sugar in the dark for 5 days. Representative seedlings were imaged (**d**), hypocotyl length (**e**) and PCK activity (**f**) were measured. Scale bar, 10 mm (**a,d**). **g,** RT-qPCR analysis of *PCK1* transcript levels relative to the constitutively expressed control gene *UBQ10* in 5-day-old *pck1* and WT seedlings grown on 0.5x MS medium with or without 2 μM PPZ in the dark. Quantitative data are mean ± s.d., *n* = 16 seedlings (**b,e**), *n* = 3 biological replicates (**c,f,g**). Different letters reflect significant differences between means (one-way ANOVA, Tukey’s test, *P <* 0.05). **h,** Anti-GFP immunoblot analysis of PCK1-YFP protein levels in 5-day-old *PCK1-YFP/pck1* seedlings grown in the dark on 0.5x MS medium containing different concentrations of PPZ. CBB: Coomassie Brilliant Blue staining of the gel blot.

### BIN2 phosphorylates PCK1 at Ser-62 and Thr-66

We performed immunoprecipitation of PCK1, using transgenic plants expressing PCK1-YFP in the *pck1* mutant (PCK1-YFP/*pck1*), followed by mass spectrometry analysis. This identified a phosphopeptide containing phosphorylation of Ser-62 and Thr-66 (Fig. 2a). Multiple sequence alignment of PCKs showed that Ser-62 and Thr-66 are conserved among various plant species (Fig. 2b; Extended Data Fig. 2). Given that Ser-62 and Thr-66 match the consensus GSK3 phosphorylation sites, we investigated whether PCK1 is a substrate of BIN2. We first expressed and purified recombinant 6×His-SUMO-PCK1 and GST-BIN2 (Extended Data Fig. 3) and performed *in vitro* kinase assays. LC-MS/MS analysis identified the same phosphopeptide containing phospho-Ser-62 and phospho-Thr-66 (Extended Data Fig. 4). To further confirm S62 and T66 are the major phosphorylation sites, we mutated Ser-62 and Thr-66 to alanine (PCK1^S62A/T66A^, or PCK1^AA^). We expressed and purified recombinant proteins, including 6×His-SUMO-PCK1, 6×His-SUMO-PCK1^AA^, GST-BIN2, GST-BIN2^K69R^ (a kinase-dead form of BIN2)^[6]^, and GST protein (Extended Data Fig. 3). After *in vitro* kinase assays, the phosphorylation status of PCK1 was analyzed by Phos-tag gel immunoblot. The results show that BIN2 phosphorylates PCK1 but not PCK1^AA^ (Fig. 2c). Next, we transiently expressed PCK1-YFP and PCK1^AA^-YFP alone or together with YFP or BIN2-YFP in *N. benthamiana* leaves. We confirmed the expression of these proteins by confocal microscopy (Extended Data Fig. 5a) and immunoblotting (Extended Data Fig. 5b) and then analyzed the phosphorylation of PCK1 proteins using Phos-tag gel blotting. Consistent with the *in vitro* observation, co-expression with BIN2-YFP caused phosphorylation of PCK1-YFP but not PCK1^AA^-YFP in *N. benthamiana* leaves (Fig. 2d). These *in vitro* and *in vivo* experiments demonstrate that BIN2 phosphorylates PCK1 at Ser-62 and Thr-66 residues.

**Fig. 2:**
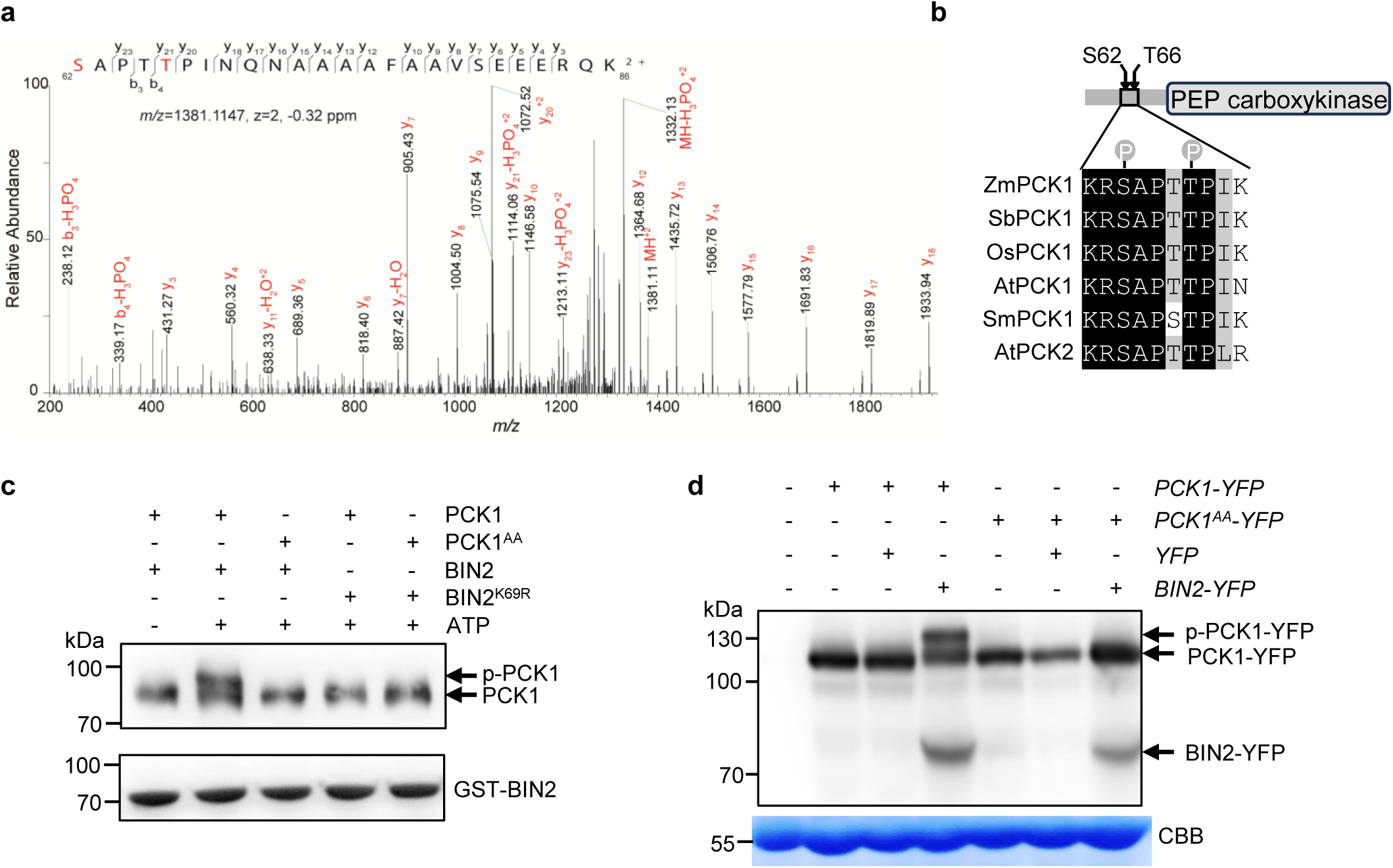
BIN2 phosphorylates PCK1 at conserved Ser-62 and Thr-66. **a,** MS/MS spectra of phosphopeptide 62-KRSAPTTPINQNAAAAFAAVSEEER-86 of PCK1-YFP protein immunoprecipitated from 2-week-old PCK1-YFP*/pck1* transgenic plants, identifying phosphorylation at Ser-62 and Thr-66 (highlighted in red). **b,** The sequence around Ser-62 and Thr-66 of PCK1 is conserved in PCKs of *Zea mays* (Zm), *Sorghum bicolor* (Sb), *Oryza sativa (*Os), *Arabidopsis thaliana* (At), and *Selaginella moellendorffii (*Sm). **c,** BIN2 phosphorylates PCK1 *in vitro*. The recombinant 6×His-SUMO-PCK1 and 6xHis-SUMO-PCK1^AA^ proteins were incubated with GST-BIN2 or kinase-inactive mutant GST-BIN2^K69R^ protein and adenosine triphosphate (ATP), then analyzed by Phos-tag gel blot probed with anti-his-tag antibody. p-PCK1: phosphorylated PCK1. The GST-BIN2 proteins were analyzed by SDS-PAGE and anti-GST immunoblotting. **d,** BIN2 phosphorylates PCK1 *in vivo*. The indicated proteins were transiently co-expressed in *N. benthamiana* leaves and analyzed using Phos-tag gel blot probed with an anti-GFP antibody. CBB: Coomassie Brilliant Blue staining of the gel blot.

### BIN2 physically interacts with PCK1

PCK1 was found to be in proximity with BIN2 *in vivo* based on a proximity-labeling mass spectrometry study^9^. To further confirm the *in vivo* association between BIN2 *and* PCK1, we first performed a co-immunoprecipitation (co-IP) assay using Arabidopsis transformed with *pBIN2:BIN2-GFP* and *35S:GFP*. Total protein extracts were immunoprecipitated with anti-GFP beads, and the precipitated proteins were analyzed by immunoblotting with anti-PCK1 antibody^52^. The results showed co-immunoprecipitation of PCK1 by BIN2-GFP but not by GFP alone (Fig. 3a). The *in vivo* interactions were further confirmed by bi-molecular fluorescence complementation assays (Extended Data Fig. 6).

**Fig. 3:**
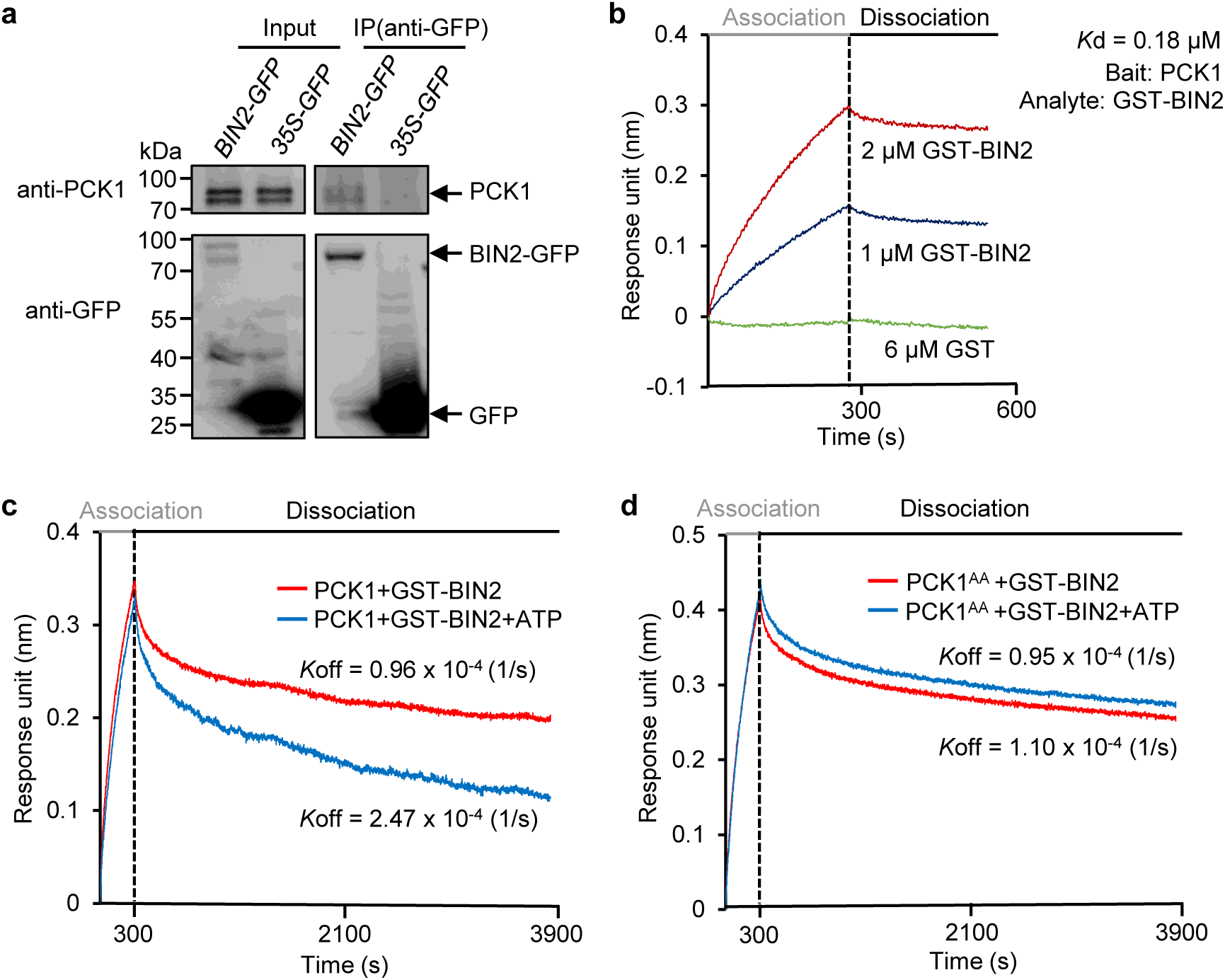
BIN2 interacts with PCK1. **a,** PCK1 co-immunoprecipitated with BIN2 in Arabidopsis. Extracts (input) from 4-week-old *pBIN2:BIN2-GFP* and *35S-GFP* transgenic plants were immunoprecipitated (IP) using anti-GFP beads and analyzed by anti-PCK1 and anti-GFP antibodies. **b,** Bio-layer interferometry (BLI) assay of the kinetics of interaction between PCK1 and BIN2. Recombinant 6×His-SUMO-PCK1 was immobilized on probes and incubated with 1 µM or 2 µM GST-BIN2 (association) followed by incubating with 1×Phosphate-buffered saline (PBS) buffer (dissociation). *K*d was determined by the *K*_off_/*K*_on_ ratio. The vertical dashed line indicates the boundary between binding and dissociation processes. **c,d,** BLI assay of the interactions between 6×His-SUMO-PCK1 or 6×His-SUMO-PCK1^AA^ and GST-BIN2 in the presence or absence of adenosine triphosphate (ATP). The experiments were repeated twice with similar results.

Next, we performed Bio-layer interferometry (BLI) assays to test the direct interaction and examine the binding kinetics between PCK1 and BIN2 using the recombinant 6×His-SUMO-PCK1, 6×His-SUMO-PCK1^AA^, and GST-BIN2 (Extended Data Fig. 3). The BLI assay revealed that PCK1 directly binds to BIN2 with a dissociation constant (*K*d) of 0.18 μM (Fig. 3b). No binding was detected between PCK1 and GST. When the BLI assay was performed in the presence of ATP in the reaction buffer, the on-rate of PCK1-BIN2 association was similar but the dissociation rate (*K*_off_) was increased by about 1.5-fold compared to the absence of ATP (Fig. 3c). ATP had no effect on the dissociation between BIN2 and PCK1^AA^ (Fig. 3d). The results indicate that BIN2 phosphorylation of PCK1 on S62/T66 enhances the dissociation of PCK1 from the kinase, thereby speeding up substrate cycling. Such effects of phosphorylation on BIN2 kinase-substrate dissociation are expected and have been observed^11,53^. Together, these results demonstrate that PCK1 is a bona fide BIN2 substrate.

### BIN2 phosphorylation of S62 and T66 inhibits PCK1

After incubation with BIN2, approximately 50% of recombinant PCK1 protein was phosphorylated, leading to activity decrease by approximately 30% (Fig. 4a). Incubation with BIN2 did not cause phosphorylation or a change in enzymatic activity of PCK1^AA^, indicating that BIN2 inhibits PCK1 by phosphorylating S62/T66 (Fig. 4a). We measured PCK1 activity after transient expression and immunoprecipitation of PCK1-YFP and PCK1^AA^-YFP from *Nicotiana benthamiana*. Consistent with the *in vitro* results, BIN2 reduced PCK1 activity but had no effect on PCK1^AA^ in *Nicotiana benthamiana* (Fig. 4b). These results demonstrate that BIN2 phosphorylation of PCK1 at Ser-62 and Thr-66 inhibits PCK1 activity.

**Fig. 4:**
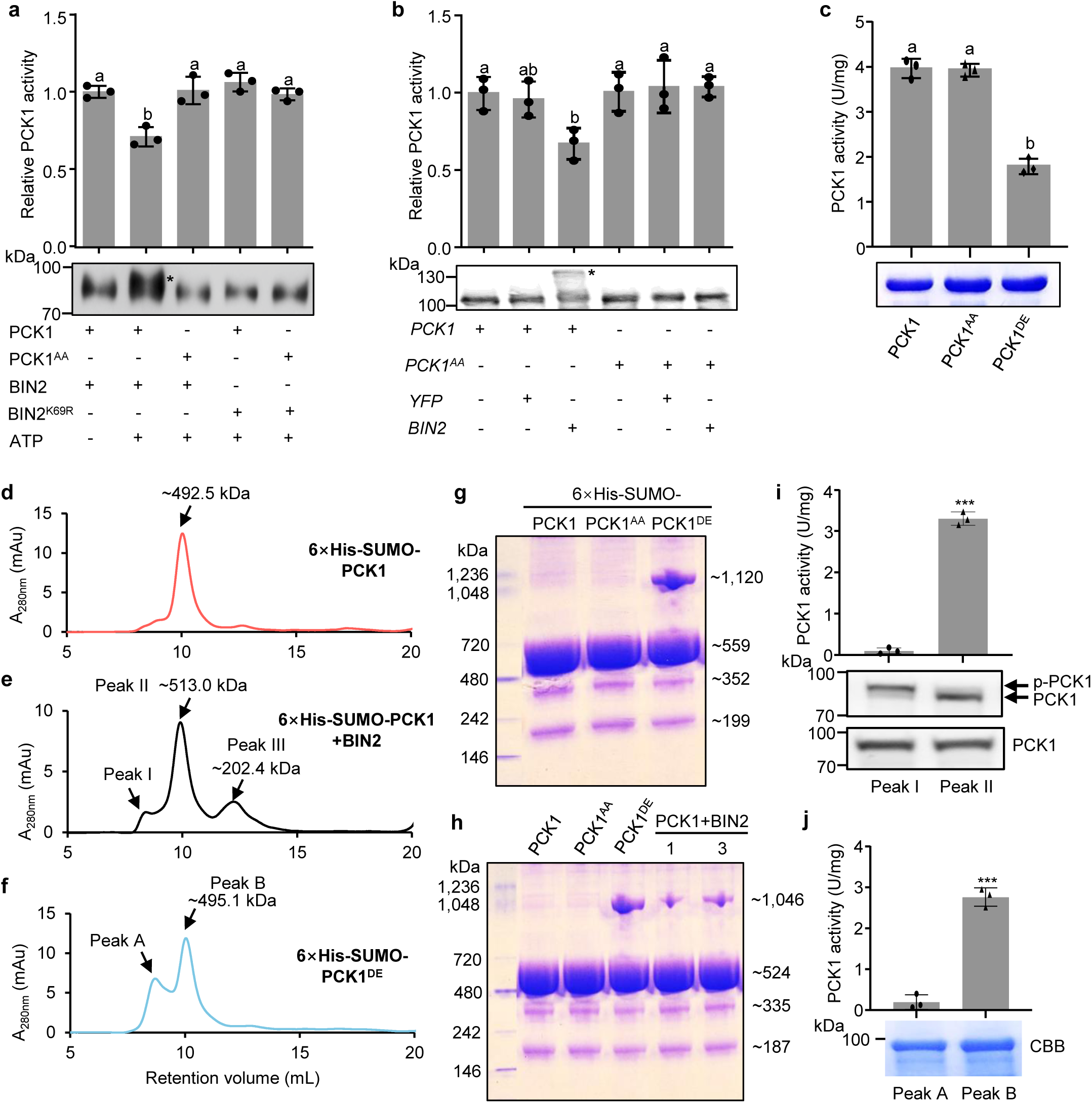
BIN2-mediated phosphorylation of S62 and T66 inhibits PCK1. **a,** Recombinant His-SUMO-PCK1 and His-SUMO-PCK1^AA^ proteins were incubated with GST-BIN2 or GST-BIN2^K69R^ in kinase reaction buffer, the decarboxylase activity was measured, and PCK1 phosphorylation was analyzed by Phos-tag gel blots probed with anti-his-tag antibody. Phosphorylated PCK1 (p-PCK1) is the shifted band marked with *. **b,** PCK1-YFP and PCK1^AA^-YFP were co-expressed with BIN2-YFP or YFP in *N. benthamiana* leaves, immunoprecipitated using anti-GFP antibody beads, and analyzed by decarboxylase assay and Phos-tag gel blot probed with anti-GFP antibody. **c,** Activities of the recombinant 6×His-SUMO-PCK1, 6×His-SUMO-PCK1^AA^, and 6×His-SUMO-PCK1^DE^ proteins. Gel images show CBB staining of SDS-PAGE analysis of the proteins. PCK activities are mean ± s.d., *n* = 3 replicates; Letters above the bars reflect significant differences between means (one-way ANOVA, Tukey’s test, *P <* 0.05). **d**-**f,** Size exclusion chromatograph (SEC) of recombinant 6×His-SUMO-PCK1 protein (**d**), 6×His-SUMO-PCK1 protein after phosphorylation by GST-BIN2 (**e**), and 6×His-SUMO-PCK1^DE^ (**f**). **g,h,** Native PAGE analyses of recombinant 6×His-SUMO-PCK1, 6×His-SUMO-PCK1^AA^, 6×His-SUMO-PCK1^DE^ (**g**), or PCK1, PCK1^AA^, PCK1^DE^, and PCK1 protein phosphorylated by GST-BIN2 *in vitro* for 1 and 3 hours (**h**). **i,** Enzymatic activity and anti-his-tag immunoblotting analysis of Phos-tag gel (upper gel panel) and SDS-PAGE gel (low panel) of the SEC fractions of peak I and peak II in (**e**). **j,** Enzymatic activity and SDS-PAGE analysis of the two peak SEC fractions of recombinant 6×His-SUMO-PCK1^DE^ shown in (**f**). PCK activities are mean ± s.d., *n =* 3 replicates; ***, *P <* 0.001, according to two-tailed Student’s *t*-test (**i,j**).

To further assess the impact of Ser-62 and Thr-66 phosphorylation on PCK1 activity, we mutated these two residues to phosphomimetic amino acids, aspartic acid and glutamic acid (PCK1^S62D/T66E^ or PCK1^DE^). We expressed and purified recombinant 6×His-SUMO-PCK1, 6×His-SUMO-PCK1^AA^, and 6×His-SUMO-PCK1^DE^ from *E. coli* and measured PCK1 activity. The results show that the PCK1 activity of 6×His-SUMO-PCK1 and 6×His-SUMO-PCK1^AA^ was comparable, at approximately 4.0 U/mg. In contrast, the activity of 6×His-SUMO-PCK1^DE^ was reduced by about 57% compared to the wild-type PCK1 (Fig. 4c). After removing the 6xHis-SUMO tag by SUMO protease digestion (Extended Data Fig. 7a), PCK1^DE^ still showed similarly reduced decarboxylase activity (Extended Data Fig. 7b), excluding the possibility of artifacts by the tags. These results indicate that the PCK1 activity is reduced by the phosphomimetic mutations.

### BIN2 phosphorylation inhibits PCK1 by promoting homo-complex formation

The Arabidopsis PCK1 and PCK2 have been shown to exist as hexamers^33^. We expressed and purified recombinant 6×His-SUMO-PCK1 and analyzed the native complex by size exclusion chromatography (SEC) using a Superdex 200 column. The unphosphorylated 6×His-SUMO-PCK1 eluted as a major peak corresponding to approximately 493 kDa (Fig. 4d; Extended Data Fig. 7c-d), consistent with the calculated size of a hexamer (522 kDa). After phosphorylation by BIN2, 6×His-SUMO-PCK1 eluted as three peaks of approximately 202, 513 kDa, and a larger size eluted near the void volume of the Superdex 200 column (peak 1) (Fig. 4e). The 6×His-SUMO-PCK1^DE^ protein fractionated as two peaks of 495 kDa and a larger size (Fig. 4f). To determine the size of the larger complex, we analyzed these proteins using blue native gel electrophoresis (BNGE) (Fig. 4g). In BNGE, 6×His-SUMO-PCK1 and 6×His-SUMO-PCK1^AA^ exhibited an apparent molecular mass of ∼559 kDa, whereas 6×His-SUMO-PCK1^DE^ showed two bands of ∼559 and 1,120 kDa, which are consistent with hexamer and dodecamer. To exclude the possibility of artifacts from the tags, we performed BNGE analysis of the full-length PCK1, PCK1^AA^, and PCK1^DE^ without the 6xHis-SUMO tag (Extended Data Fig. 7a). Both PCK1 and PCK1^AA^ showed a major band of ∼524 kDa in BNGE, which is close to the predicted size of a hexamer. Similar to 6×His-SUMO-PCK1^DE^, PCK1^DE^ exhibited two bands of ∼524 and ∼1,046 kDa. BIN2 phosphorylation generated a PCK1 complex of about 1,100 kDa (Fig. 4h), indicating that dodecamer formation is independent of the tags. BNGE also revealed minor amounts of smaller PCK1 protein complexes with measured sizes consistent with dimers and tetramers (Fig. 4g,h). These results indicate that PCK1 is predominantly a hexamer, and phosphorylation causes formation of a dodecamer, with small amounts of dimers and tetramers present at the equilibrium.

After the BIN2 phosphorylation assay, only a fraction of PCK1 shifted to the dodecamer complex. To determine whether this partial effect on complex formation is due to partial phosphorylation, we analyzed the phosphorylation status of PCK1 in the two SEC peak fractions using Phos-tag gel blot experiments. The results showed that the hexamer fraction contains mostly unphosphorylated PCK1, and the dodecamer fraction contains mostly the phosphorylated form (Fig. 4i), suggesting that phosphorylation causes formation of the dodecamer. In contrast, a significant portion of PCK1^DE^ exists in the hexamer fraction (Fig. 4f-h), suggesting that the S62D and T66E mutations only partially mimic phosphorylation in shifting the equilibrium towards the dodecamer. Enzymatic assays revealed that the dodecamer fraction of phosphorylated PCK1 exhibited a significantly reduced activity (Fig. 4i; Extended Data Fig. 7e), indicating that phosphorylation of PCK1 S62/T66 effectively inhibits PCK1 decarboxylation activity while causing the formation of a higher-order complex. Interestingly, while the dodecamer fraction of PCK1^DE^ showed significantly reduced activity like the phosphorylated PCK1, the PCK1^DE^ hexamer fraction exhibited near-normal activity (Fig. 4j), suggesting that phosphorylation *per se* is not sufficient and that formation of the dodecamer complex is required for PCK1 inactivation. These observations collectively suggest that phosphorylation inhibits PCK1 activity by promoting a quaternary structural change.

### BIN2 phosphorylation of Ser-62/Thr-66 inhibits PCK1 *in vivo*

To test whether BIN2 is required for PCK1 phosphorylation *in vivo,* we analyzed PCK1 phosphorylation in the triple loss-of-function mutant *bin2T* (*bin2-3 bil1 bil2* triple mutant) (Fig. 5a-d). When grown on 2 µM PPZ, the wild-type seedlings displayed short hypocotyls, whereas the *bin2T* mutant showed a significantly reduced sensitivity to PPZ (Fig. 5a,b). Immunoblotting shows higher levels of PCK1 phosphorylation in WT than *bin2T* (Fig. 5c,d), providing genetic evidence for the essential function of BIN2 and its homologs in phosphorylating PCK1 *in vivo*. To determine whether *in vivo* BIN2 phosphorylation of PCK1 requires Ser-62 and Thr-66, we generated transgenic plants that express the wild-type *PCK1-YFP* and the unphosphorylatable mutant *PCK1^AA^-YFP* in the *pck1* mutant background (Extended Data Fig. 8). We crossed these transgenic lines into the gain-of-function *bin2-1* background. Phos-tag gel blotting analysis detected phosphorylation of PCK1-YFP, but not PCK1^AA^-YFP in *bin2-1* (Fig. 5e). While the PCK1 expression levels are similar between the seedlings of *PCK1-YFP/bin2-1* and *PCK1^AA^-YFP /bin2-1,* the PCK decarboxylase activity of *PCK1-YFP/bin2-1* and *PCK1-YFP ^AA^/bin2-1* is 33% and 139% higher, respectively, than that of *bin2-1* (Fig. 5f). These results demonstrate that BIN2 phosphorylates Ser-62 and Thr-66 to inhibit PCK1 decarboxylase activity.

**Fig. 5:**
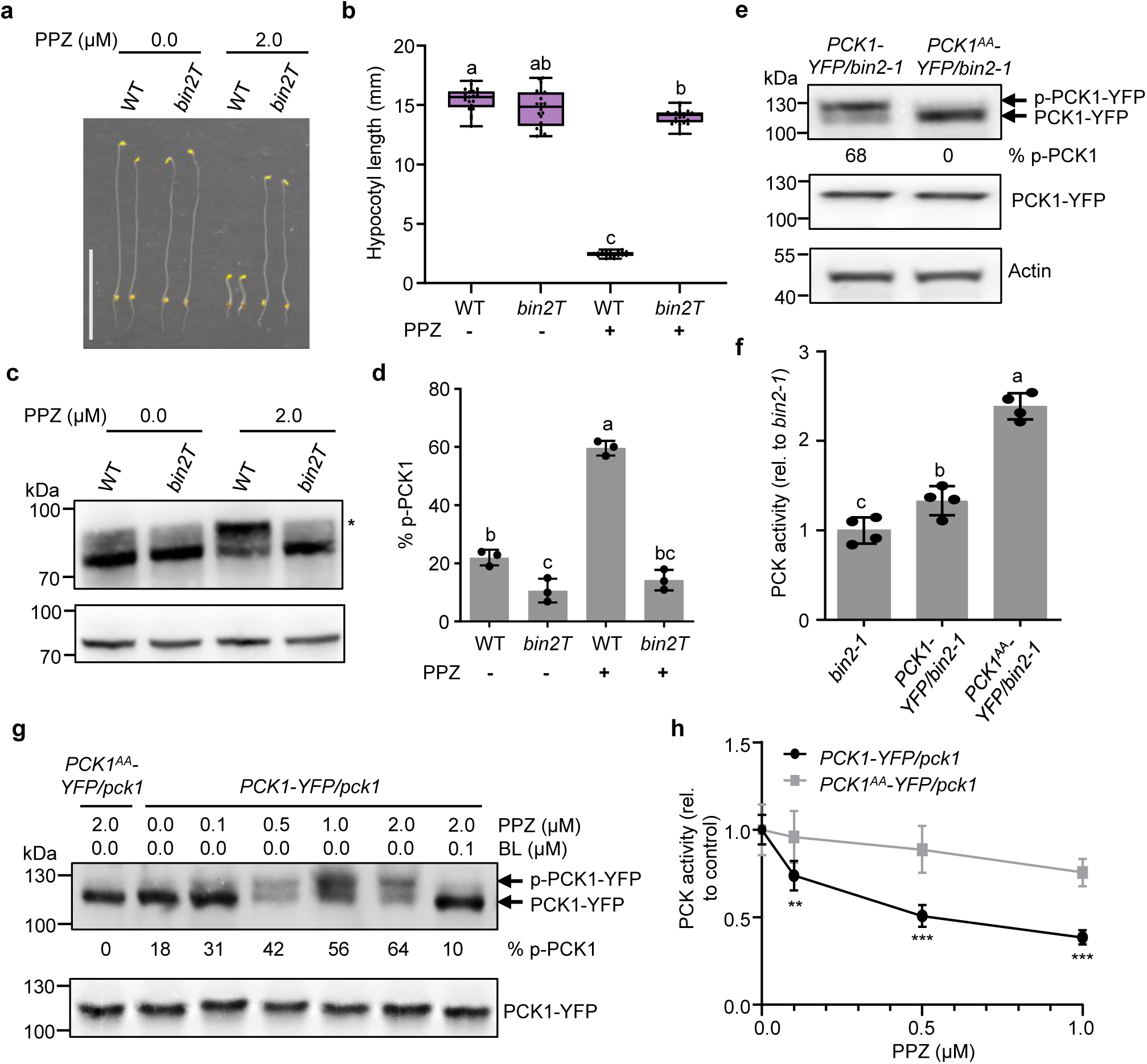
BIN2 phosphorylation of Ser-62 and Thr-66 mediates BR regulation of PCK1 activity. **a,** Representative seedlings of wild type (WT) and *bin2T* (*bin2-3 bil1 bil2* triple mutant) grown on sugar-free medium with or without 2 μM propiconazole (PPZ) in the dark for 4 days. Scale bar: 10 mm. **b,** Hypocotyl lengths of seedlings in (**a**). The central lines within the box plots represent the medians, the box represents the interquartile range (IQR), the whiskers extend to minima and maxima. *n =* 21 seedlings. **c,** Phosphorylation status of PCK1 in seedlings shown in (**a**) was analyzed by Phos-tag gel blot (upper panel) or SDS-PAGE (lower panel) probed with an anti-PCK1 antibody. Phosphorylated PCK1 (p-PCK1) is the shifted band marked with *. **d,** The % of p-PCK1 was calculated from the band intensities of three replicates of the immunoblot shown in (**c**). Data are mean ± s.d., Different letters reflect significant differences between means (one-way ANOVA, Tukey’s test, *P <* 0.05). **e,** Immunoblots of Phos-tag gel (upper panel) and SDS-PAGE (lower panels) show the phosphorylation status of PCK1-YFP in 4.5-day-old etiolated seedlings of *PCK1-YFP/bin2-1* and *PCK1^AA^-YFP/bin2-1* grown on 0.5x MS medium in the dark. Anti-actin immunoblotting was used as an internal control. **f,** Relative PCK decarboxylase activity of *bin2-1, PCK1-YFP/bin2-1* and *PCK1^AA^-YFP/bin2-1*, Data are mean ± s.d., *n =* 4 biological replicates. Different letters reflect significant differences between means (one-way ANOVA, Tukey’s test, *P <* 0.05). **g,** The *PCK1^AA^-YFP/pck1* and *PCK1-YFP/pck1* seedlings were grown in the dark on 0.5x MS medium containing the indicated concentrations of PPZ and brassinolide (BL). The proteins were analyzed by Phos-tag (upper panel) or non-Phos-tag gel (lower) and probed with an anti-GFP antibody. The % of p-PCK1-YFP was calculated from the band intensities. **h,** Relative PCK decarboxylation activity of samples described in (**g**). Data are mean ± s.d., *n* = 3 biological replicates. Significant differences from seedlings grown without PPZ: **, *P <* 0.01, ***, *P <* 0.001 (one-way ANOVA, Tukey’s test, *P <* 0.05).

### BR activates PCK1 by inhibiting Ser62/Thr66 phosphorylation

To confirm BR regulation of PCK1 phosphorylation *in vivo*, we analyzed PCK1 phosphorylation status in two BR biosynthetic mutants, *det2* and *dwf4,* grown on medium supplemented with 0, 4, and 10 nM brassinolide. The growth of *det2* and *dwf4* was gradually rescued by the increasing concentrations of brassinolide (Extended Data Fig. 9a,b). Phosphorylation levels of PCK1 are higher in *det2* and *dwf4* than WT, but reduced by brassinolide treatment to a similar level as WT (Extended Data Fig. 9c,d), demonstrating that BR regulates PCK1 phosphorylation. In the root tip, BR levels are low in the meristem and high in the elongation zone (Extended Data Fig. 9e)^54^. We separately harvested tissues of the meristem (∼0.5 mm) and elongation zone (0.5-1 mm) and analyzed PCK1 phosphorylation by Phos-tag gel blotting. The results show that PCK1 is more phosphorylated in the meristem than in the elongation zone (Extended Data Fig. 9f-g). Together, these results confirm the tight correlation between BR level and PCK1 dephosphorylation *in vivo*.

To demonstrate the causal relationship between BIN2 phosphorylation of PCK1 Ser-62/Thr-66 and BR regulation of PCK1 activity, we grew the transgenic seedlings expressing PCK1-YFP and the unphosphorylatable mutant PCK1^AA^-YFP on medium supplemented with a range of concentrations of PPZ and analyzed the phosphorylation status of PCK1-YFP using Phos-tag gel immunoblot. The results showed that PPZ increased the phosphorylation of PCK1-YFP, but had no effect on the phosphorylation status of PCK1^AA^-YFP (Fig. 5g). The effect of PPZ was abolished by exogenous co-application of 0.1 μM brassinolide (Fig. 5g). PCK1 protein levels of both *PCK1-YFP/pck1 and PCK1^AA^-YFP/pck1* transgenic seedlings were not changed by PPZ treatment (Extended Data Fig. 8b). These results suggest that Ser-62 and Thr-66 are essential for BR regulation of PCK1 phosphorylation status in Arabidopsis. Consistent with the effects on phosphorylation, PPZ reduced the PCK activity in PCK1-YFP/*pck1* seedlings but had little effect on PCK1^AA^ (Fig. 5h). These results demonstrate that BR activates PCK1 by inhibiting BIN2 phosphorylation of Ser62 and Thr66.

### BR activation of PCK1 contributes to seedling growth

To evaluate the functional significance of PCK1 phosphorylation in plant development, we analyzed the growth phenotypes of transgenic lines expressing PCK1-YFP, PCK1^AA^-YFP, and PCK1^DE^-YFP. Three independent transgenic lines of each transgene that expressed similar PCK1 levels to endogenous PCK1 levels of wild-type seedlings were selected for measurement of PCK1 activity (Fig. 6a; Extended Data Fig. 8a). When grown on 0.5x MS sugar-free medium in the dark, the *PCK1-YFP/pck1* seedling showed similar phenotypes as wild type (Fig. 6a,b). The hypocotyls of *PCK1^AA^-YFP/pck1* plants were about 32.4% longer than *PCK1-YFP/pck1*, while those of *PCK1^DE^-YFP/pck1* lines were 16.0% shorter than *PCK1-YFP/pck1* (Fig. 6a,b). The PCK activity of *PCK1^AA^-YFP/pck1* seedlings was 27.7% higher than *PCK1-YFP/pck1* seedlings, whereas *PCK1^DE^-YFP/pck1* showed 21.8% lower activity than *PCK1-YFP/pck1* seedlings (Fig. 6c). Thus, the hypocotyl lengths positively correlated with the PCK activity in these transgenic lines. To test whether the different growth phenotypes were due to different sugar levels, we grew these plants on media containing sucrose. Indeed, the hypocotyl lengths of these transgenic lines were similar when grown on a sugar-containing medium (Extended Data Fig. 10a,b).

**Fig. 6:**
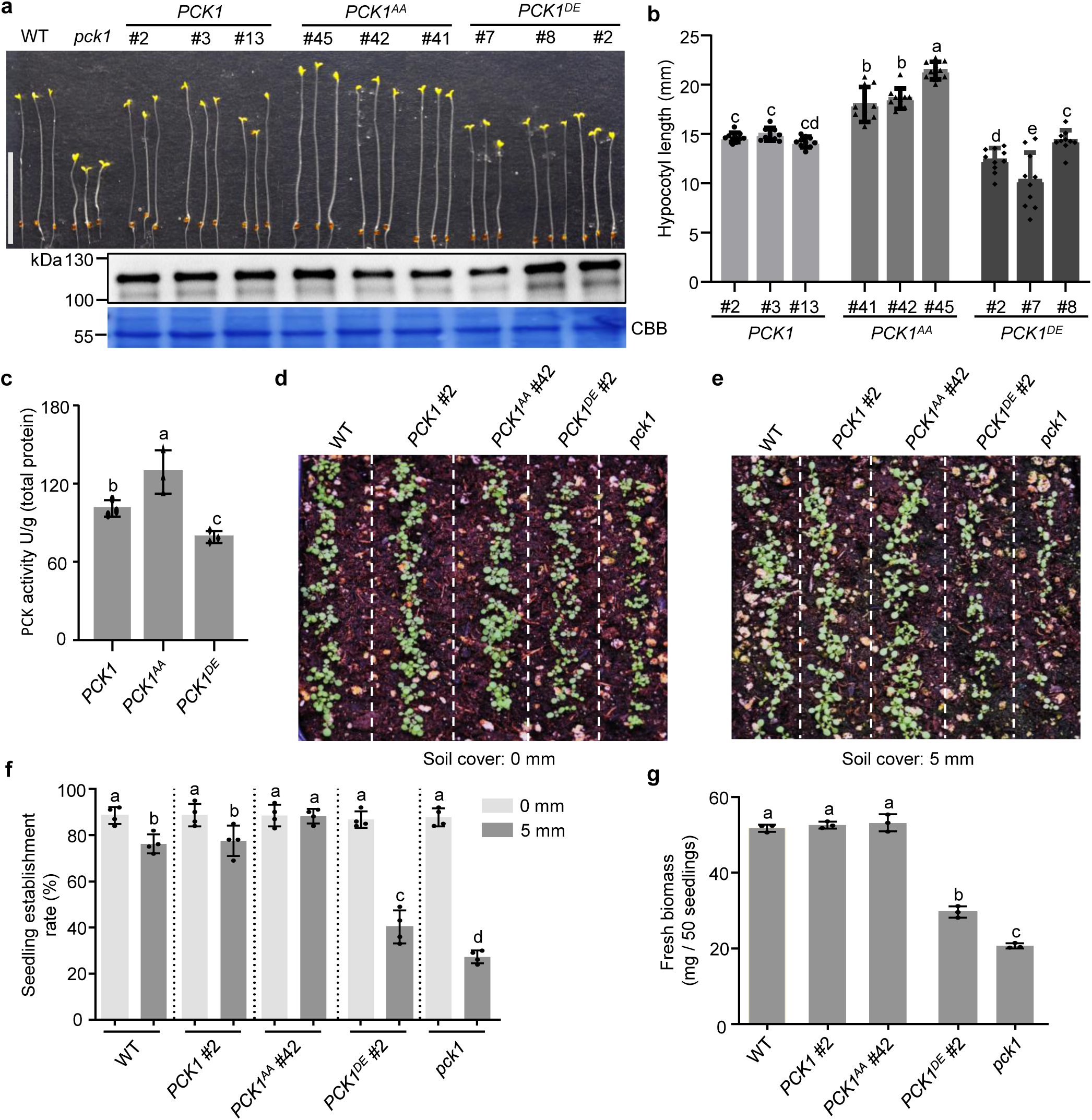
Activation of PCK1 by dephosphorylation enhances seedling growth and emergence from soil. **a**-**c,** Seedlings of wild-type (WT), *pck1*, *PCK1-YFP/pck1* (*PCK1*), *PCK1^AA^-YFP/pck1* (*PCK1^AA^*), and *PCK1^DE^-YFP/pck1* (*PCK1^DE^*) transgenic lines were grown in the dark for 5 days. Three representative seedlings were imaged (**a**, upper panel; scale bar: 10 mm), PCK1-YFP protein levels were analyzed by immunoblot (**a**, lower panels), hypocotyl lengths were measured from 10 seedlings (**b**), and the PCK activities were measured for the three transgenic lines (**c**). **d**-**g,** Seeds of the indicated genotypes were sown on the surface of soil (**d**) or covered by 5 mm of soil (**e**), and grown under light for 7 days. **f,** Seedling establishment rate is the ratio between the number of seedlings and the number of seeds sown. **g,** Fresh weight of 50 seedlings shown in (**d**). Data are mean ± s.d., *n* = 10 seedlings (**b**), *n* = 3 independent transgenic lines (**c**), *n* = 4 biological replicates (**f**), *n* = 3 of 50 pooled seedlings (**g**); Different letters reflect significant differences between means (one-way ANOVA, Tukey’s test, *P <* 0.05).

We next tested whether PCK1 phosphorylation influences seed germination and seedling establishment under different growth conditions. We sowed the seeds of WT, *pck1*, *PCK1-YFP/pck1, PCK1^AA^-YFP/pck1, and PCK1^DE^-YFP/pck1* on agar-solidified medium or on the surface of soil and put them under light or in the dark. The germination rate of all genotypes on medium was about 95% under both light and dark conditions (Extended Data Fig. 10c). The germination rate of all genotypes on soil was about 90% under both light and dark conditions (Extended Data Fig. 10d). We then sowed 140 seeds of each genotype on the surface (Fig. 6d) or at a soil depth of 5 mm in each pot of soil (Fig. 6e). A week later, about 90% of the seeds of all genotypes sown on the surface germinated (Fig. 6d,f). Covering the seeds with 5 mm of soil reduced the germination/emergence rate of WT and *PCK1-YFP/pck1* by about 14%, of the *pck1* mutant by 69%, and the *PCK1^DE^-YFP/pck1* by 55% (Fig. 6e,f). In contrast, the emergence rate of *PCK1^AA^-YFP/pck1* is about 88% both on the soil surface and at 5 mm depth, showing no significant decrease by soil coverage (Fig. 6f). We pulled out 50 seedlings from each sample and measured their fresh weight. The WT, *PCK1-YFP/pck1*, and *PCK1^AA^-YFP/pck1* seedlings grown on soil surface accumulated similar biomass (Fig. 6g). The fresh weight of *PCK1^DE^-YFP/pck1* and *pck1* seedlings were reduced significantly to about 40% and 24% of WT when grown on soil surface, respectively (Fig. 6g). Together, these observations demonstrate that phosphorylation by BIN2 inhibits PCK1 activity, reduces sugar availability, and thus delays seedling growth and impedes seedling establishment in soil. In contrast, BR-induced PCK1 dephosphorylation and activation increases sugar synthesis from lipid storage and is crucial for seedling establishment in natural soil conditions.

### BIN2-mediated phosphorylation inhibits PCK1 in maize and sorghum

The BIN2 phosphorylation sites, Ser-62 and Thr-66 of Arabidopsis PCK1, exist within a short sequence, RSAPTTP, which is highly conserved among most PCK homologues in different plant species (Fig. 2b; Extended Data Fig. 2). Phosphorylation of these conserved residues has been observed in association with PCK inactivation in maize (Ser-55 and Thr-59)^47^. To investigate whether BIN2 regulation of PCK1 is conserved across other plant species, especially in C4 crops, we treated maize and sorghum leaves with brassinolide or bikinin and analyzed PCK1 phosphorylation and enzymatic activity. In both maize (Fig. 7a) and sorghum (Fig. 7b) grown under an 11h-light/13h-dark photoperiod, the majority of PCK1 was dephosphorylated at the end of the light period. After 4.5 and 9 hours in the dark period, phosphorylation levels increased, and PCK enzymatic activity decreased (Fig. 7a,b). Brassinolide and bikinin treatments caused PCK1 dephosphorylation and increased PCK1 activity in both maize and sorghum (Fig. 7a,b). These results show that the phosphorylation and activity of PCK1 are diurnally regulated by BR signaling via BIN2 phosphorylation in the leaves of maize and sorghum, uncovering a potential link between BR signaling and C4 photosynthesis.

**Fig. 7:**
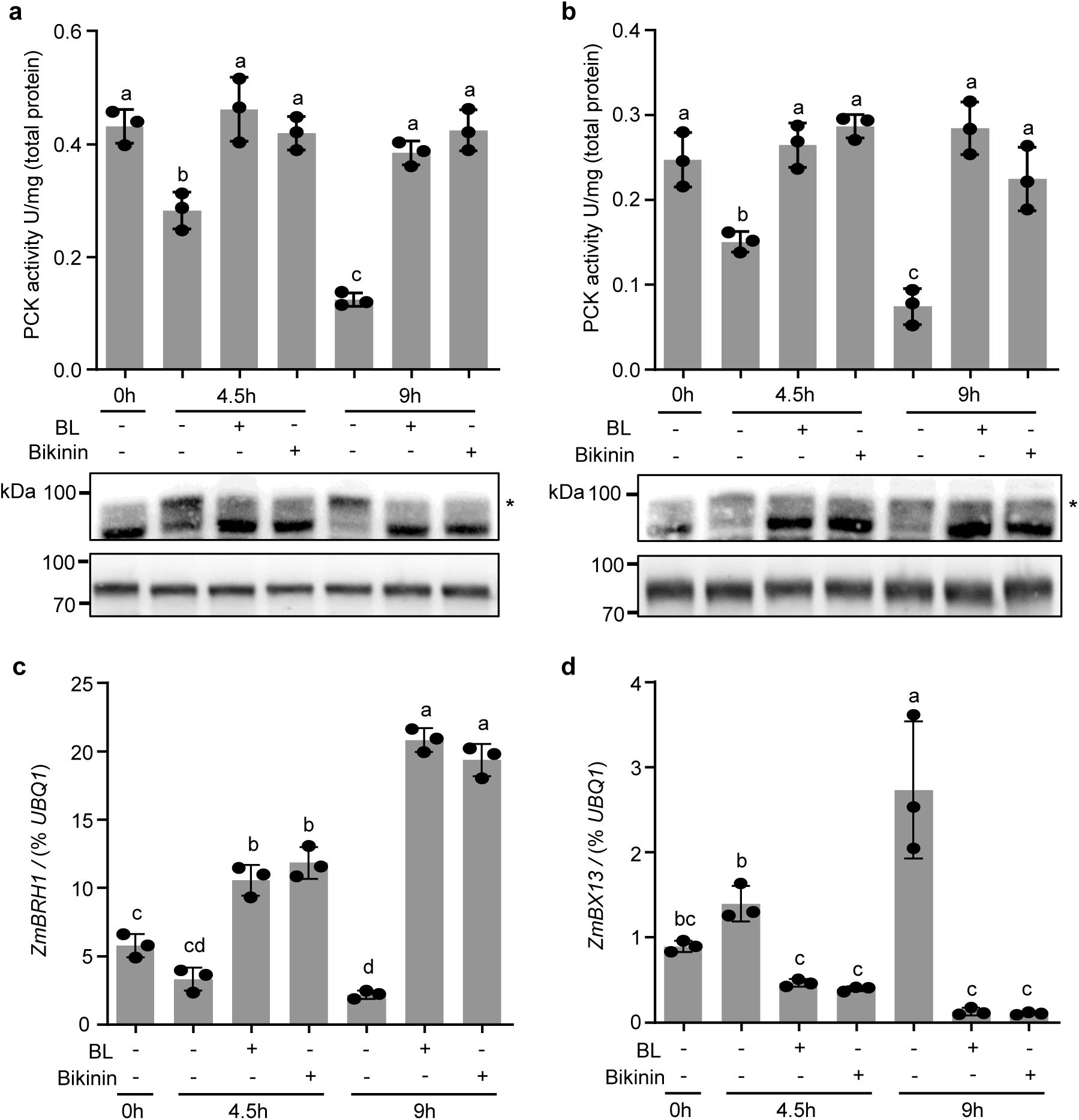
BRs regulate PCK1 through BIN2 in maize and sorghum. Quantification of PCK activity and immunoblotting of PCK1 in maize (**a**) and sorghum (**b**). Maize and sorghum plants were grown under 11h-light/13h-dark photoperiod for 5 and 6 weeks, respectively. Tissues were harvested from the middle region of leaves at the end of the day and treated with mock, 200 nM BL, or 50 µM bikinin solution in the dark for 4.5 and 9 hours. The decarboxylase activity was measured in total protein extracts, and PCK1 protein was analyzed by Phos-tag gel blots (upper panels) or SDS-PAGE (lower panels) probed with an anti-PCK1 antibody. The shifted band of phosphorylated PCK1 is marked with *. **c,d,** RT-qPCR analysis of transcript levels of *ZmBRH1* and *ZmBX13* relative to the constitutively expressed control gene *ZmUBQ1* in the maize leaves described in (**a**). Data are mean ± s.d. of 3 biological replicates. Letters above the bars reflect significant differences between means (one-way ANOVA, Tukey’s test, *P <* 0.05).

The increased PCK1 phosphorylation at night and its reversal by BR treatment suggest that BR levels are high during the day and low at night. Indeed, previous studies have shown such diurnal changes of BR levels in Arabidopsis^55,56^. We further analyzed the transcript levels of BR-responsive genes in maize by quantitative reverse transcription PCR (qRT-PCR) (Fig. 7c,d). The expression of *ZmBRH1* (Brassinosteroid-Responsive Ring-H2, Zm00001d045568), a BR-induced gene^57^, decreased in the dark and increased by BR and bikinin treatments (Fig. 7c)., whereas the expression of *ZmBX13* (Benzoxazinone synthesis13, Zm00001d007718), a BR-repressed gene, increased in the dark and was repressed by BR and bikinin (Fig. 7d). These results suggest that BR levels are high during the day and low at night in maize, similar to the observations in Arabidopsis^55,56^. Together, these results suggest that diurnal changes in BR levels contribute to the phosphorylation/inhibition of PCK1 at night and dephosphorylation/activation during the day, thereby promoting photosynthetic metabolism.

## Discussion

The interplay between sugar and BR is the core mechanism of plant growth regulation. BR induces growth by increasing cell expansion and cell wall synthesis, which consume sugars as energy sources and substrates for cell wall synthesis. Sugar is essential for viability and is monitored by sugar-sensing and signaling mechanisms to maintain homeostasis^24^. Recent studies have elucidated how sugar signaling modulates the level of key BR signaling components to ensure sugar regulation of BR-dependent growth^23–25,58^. Here, we demonstrate that BR promotes sugar synthesis. We uncover a mechanism by which GSK3/BIN2-mediated phosphorylation inhibits PCK1 catalytic activity, thereby mediating BR regulation of gluconeogenesis during seedling establishment in Arabidopsis. Our results in maize and sorghum further indicate that BR mediates diurnal regulation of PCK1 activity in C4 photosynthesis (Fig. 8).

**Fig. 8:**
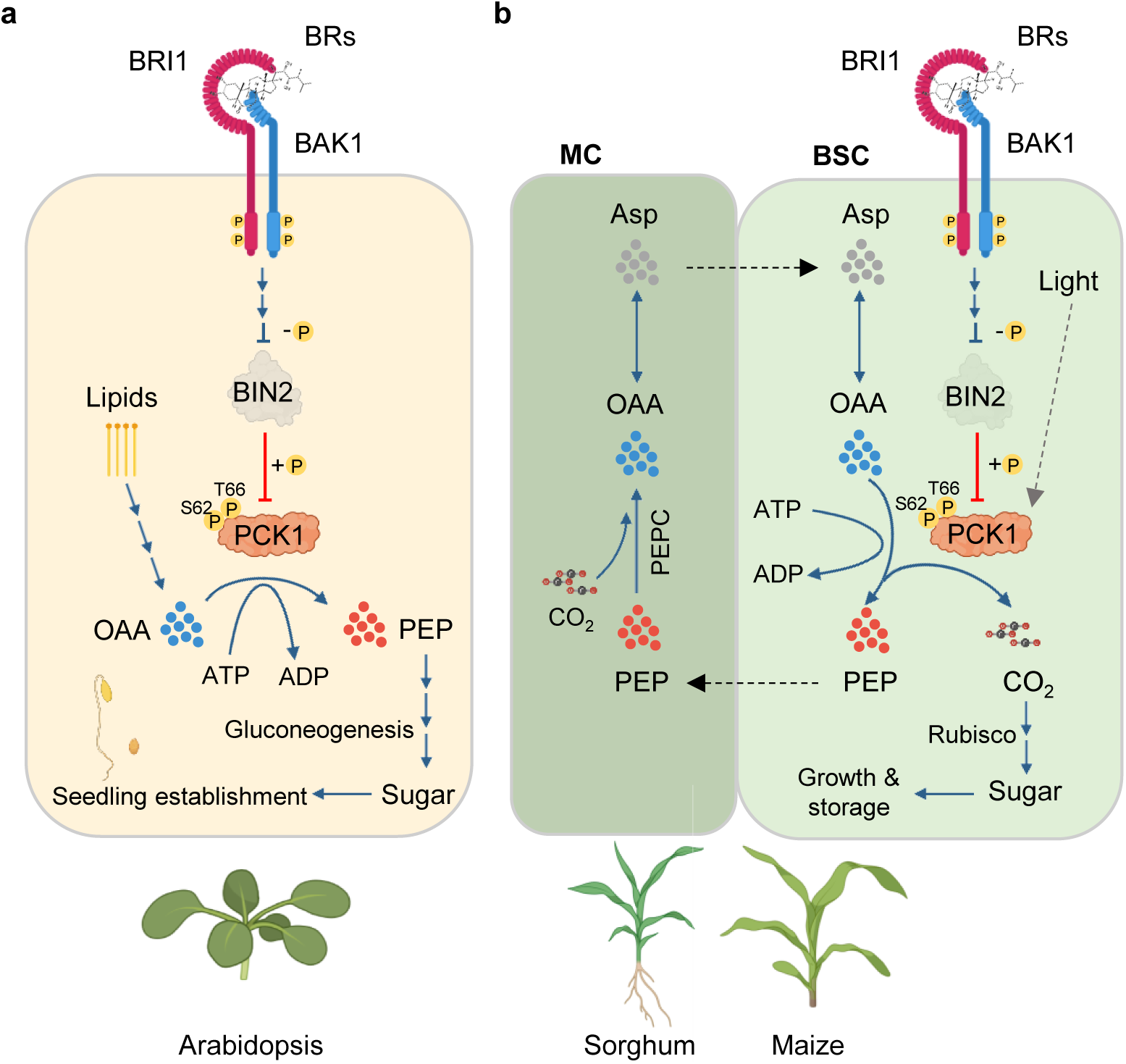
Summary of the roles of the BIN2-PCK1 regulatory module in Arabidopsis and maize/sorghum. **a,** BRs activate PCK1 by inhibiting BIN2 phosphorylation, promoting the conversion of lipids into sugar for Arabidopsis seedling establishment. **b,** BRs and light regulate PCK in bundle sheath cells (BSC) for photosynthesis in maize and sorghum. CO_2_ is fixed by PEP carboxylase (PEPC) into OAA in mesophyll cells (MC). OAA is then converted to Aspartate (Asp) and transported to BSC, where OAA is decarboxylated by PCK, thus releasing CO_2_ for photosynthesis. PEP returns to the MC for the next round of the C4 cycle.

Previous proteomic studies have shown BR regulation of PCK1 phosphorylaiton^27,28^, and our recent study identified PCK1 as a BIN2 kinase substrate *in vivo*^9,27^. Following these studies, we demonstrate that PCK1 activity is decreased in BR-deficient plants and in gain-of-function *bin2-1* mutants, but increased by treatments with BR and BIN2 inhibitor. We confirm that PCK1 phosphorylation levels correlated with BR-deficiency and BIN2 activity *in vivo*, *In vitro,* BIN2 directly interacts with PCK1 (*K*d = 0.18 µM) and phosphorylates PCK1 at Ser-62 and Thr-66, the same residues dephosphorylated *in vivo* in response to BR treatment. As expected for kinase-substrate interaction, phosphorylation increases the dissociation rate of BIN2-PCK1 interaction. The phosphoblocking mutation PCK1^AA^ abolishes BIN2 phosphorylation *in vitro*, confers BR-independent constitutive PCK1 activity *in vivo*, and promotes seedling growth on sugar-free media in the dark. Conversely, the phosphomimetic mutation PCK1^DE^ reduced PCK1 activity and compromised seedling growth, particularly seedling establishment from under the soil. These results demonstrate conclusively that BIN2 phosphorylates PCK1 at S62 and T66 to inhibit its catalytic activity, and BR activates PCK1 by inhibiting BIN2 phosphorylation, thereby promoting gluconeogenesis and post-germination seedling establishment.

We further show that phosphorylation at Ser-62 and Thr-66 inhibits PCK1 by changing its quaternary conformation. Plant PCKs are known to form homo-hexamer complexes^33,59^. Our SEC and native gel electrophoresis analyses showed that the unphosphorylated PCK1 forms an active hexamer and the phosphorylated PCK1 exists mostly in a dodecamer that exhibits nearly complete loss of enzymatic activity. Interestingly, only a small fraction of the PCK1^DE^ protein purified from *E. coli* forms dodecamer with reduced activity, while a major fraction remains hexamer with normal activity. While the S62D and T66E mutations seem to have a weaker effect than phosphorylation on dodecamer formation, the results suggest that PCK1 is inhibited by phosphorylation-induced dodecamer formation. Our study thus uncovers a novel mechanism for posttranslational regulation of PCK catalytic activity via phosphorylation-induced quaternary complex formation. The structural basis of this regulatory mechanism will be an interesting topic for future investigation.

The sequence flanking Ser-62 and Thr-66, *i.e*., KRSAPTTP, is highly conserved in PCKs across multicellular plants (Fig. 2 and Extended Data Fig. 2)^49^. This sequence is present in both PCK1 and PCK2 in Arabidopsis and duplicated in one of the PCK isoforms in maize. Therefore, GSK3-mediated regulation of PCK is likely used broadly across diverse organisms for metabolic regulation. A previous study showed that site-directed mutagenesis of either Ser-62, Thr-65, Thr-66, or a triple mutation of all three residues in PCK from multiple organisms did not support the idea that phosphorylation inhibits PCK^[49]^. It is possible that PCK inactivation requires the phosphorylation of both Ser-62 and Thr-66, as well as the unphosphorylated Thr-65. Future structural studies will provide further insight into the mechanism of PCK regulation by phosphorylation and conformational changes. Our BR and bikinin treatments of maize and sorghum indicate that the mechanism of BR activating PCK through inhibiting BIN2 phosphorylation is conserved in monocots.

PCK plays a crucial role in photosynthetic CO_2_ fixation in C4 and CAM plants^42^. Regulation of PCK activity contributes to light-harvesting plasticity and improves photosynthesis efficiency under natural diurnal conditions and particularly under stress conditions in C4 plants^42^. Studies in several plant species have demonstrated that PCK is phosphorylated at the N-terminal region, including residues corresponding to S62 and T66 in Arabidopsis PCK1, and that phosphorylation is correlated with a decrease in PCK decarboxylation activity^44,47,48,60,61^. In several C4 organisms, PCK is phosphorylated and inhibited under dark (nighttime) and shade conditions, but is dephosphorylated and active under illuminated conditions to catalyze the decarboxylation of OAA, releasing CO_2_ for fixation by Rubisco in bundle sheath cell^44,45^. Our study demonstrates that PCK is phosphorylated and inhibited in maize and sorghum in the dark, consistent with previous reports that shade inhibits PCK activity in maize^45^. We further show that BR and bikinin treatment causes PCK dephosphorylation and activation in maize and sorghum. Our analysis of BR-responsive genes indicates that BR levels are high during the day and low at night in maize, consistent with previous findings of diurnal fluctuations of BR levels in Arabidopsis^55,56^. Our study thus uncovers a role for BR signaling in the diurnal regulation of PCK activity and photosynthetic carbon fixation. Together, our work in Arabidopsis and C4 plants uncovers a GSK3/BIN2-PCK pathway by which BR promotes sugar production through gluconeogenesis and photosynthesis, thereby ensuring a sufficient sugar supply to meet the growth demands generated by other BR signaling branches. As BR enhances photosynthesis and biomass accumulation^62–64^, the molecular link between BR signaling and C4 photosynthesis provides a specific target for improving plant productivity by agrichemical and genetic approaches.

## Methods

### Plant materials and growth conditions

The *Arabidopsis thaliana* ecotypes Columbia-0 (Col-0) and Wassilewskija (WS) were used in this study. *pck1* (*pck1-2*, SALK_032133C), *bzr1-1D*, *bin2-1*, *det2-1*, *dwf4-102* (SALK_020761)*, 35S:GFP*, *and 35S:BIN2-GFP* transgenic lines are in the Col-0 background. *bin2T* is in the WS background. The *35S:PCK1-YFP*/*bin2-1* and *35S:PCK1^AA^-YFP/bin2-1* were generated by genetic crossing between the *35S:PCK1-YFP* and *35S:PCK1^AA^-YFP* with *bin2-1*. A homozygous line was identified in the F2 by screening with Basta resistance and phenotype of *bin2-1*, and propagated to F3 for experiments. For cultivation, seeds were surface-sterilized in chlorine gas for 3 h and sown on 120-mm square polystyrene petri dishes (Greiner Bio-One) containing 0.5x MS salts (Murashige and Skoog, Duchefa), with 0.8% (w/v) agar (Type M, Sigma, Steinheim). Plates were kept in the dark at 4°C for 3 days before being transferred to a growth chamber (CLF Plant Climatics) with constant darkness at 22°C in a vertical orientation, unless indicated otherwise. *Nicotiana benthamiana* plants were grown in a growth chamber at 22°C under 16 h light/8 h dark cycle. Five to six-week-old *Nicotiana benthamiana* plants were used for *Agrobacterium*-mediated infiltration.

To test seed germination and seedling growth in soil, seeds were counted and sown on the surface of soil, either uncovered or covered by 5 mm of additional soil. The pots of soil with seeds were kept for 3 days at 4°C before being placed in a growth chamber under 16-h light and 8-h dark photoperiod. Maize (*Zea mays L.*) seedlings (inbred B73) and sorghum (*Sorghum bicolor*) seedlings (inbred Tx430) were grown in a greenhouse under 11h-light/13h-dark photoperiod at 25°C.

For brassinolide and bikinin treatment of the 5-week-old maize and 6-week-old sorghum leaves, the middle region of leaves (around 50-100 mm from the tip) were cut into small pieces (around 10 mm × 20 mm) and soaked in liquid 0.5x MS medium containing 200 nM brassinolide or 50 µM bikinin under dark for 4.5 and 9 hours.

### Plasmid construction for stable *A. thaliana* transformation

To generate the construct *35S:PCK1-YFP, AtPCK1* coding sequence amplified from cDNA (All PCR primer sequences used in this study are listed in Supplementary Table 1) was cloned into a pENTR/D-TOPO entry vector through the BP reaction (Fisher Scientific). Subsequently, the *PCK1* coding sequence was transferred from the entry vector to the gateway-compatible vector pEarleyGate 101^[65]^ through the LR reaction (Fisher Scientific). To make *35S:PCK1^AA^-YFP* and *35S:PCK1^DE^-YFP*, the pENTR/D-TOPO entry vector containing *AtPCK1* coding sequence was mutated using the Q5 Site-Directed Mutagenesis Kit (NEB, E0552). The mutated PCK1 coding sequences were transferred from the pENTR/D-TOPO entry vector to the pEarleyGate 101 through the LR reaction (Fisher Scientific). All constructed plasmids were sequenced to ensure accuracy. The constructs were transformed into the *Agrobacterium tumefaciens* strain GV3101 followed by the stable transformation of *pck1* mutant plants using the floral-dip method^66^.

### Hypocotyl length measurement

Seedlings were photographed with a digital camera (Nikon D600), and hypocotyl length of seedlings was measured with ImageJ^67^.

### PCK1 phylogenetic analysis

The PCK1 amino acid sequences were retrieved from the National Center for Biotechnology Information (NCBI) database (http://blast.ncbi.nlm.nih.gov/Blast.cgi). MEGA 6.0 software was used for phylogenetic analysis using the neighbor-joining method^68^.

### RT-qPCR

Total mRNA was extracted using the RNeasy Plant Mini Kit (Qiagen) following the manufacturer’s protocol. Reverse transcription was carried out on 0.5 μg total RNA after DNase I-digestion (New England Biolabs) using the SuperScript III kit (Fisher Scientific) and oligo(dT) primers. RT-qPCR was performed with first-strand cDNAs using GoTaq qPCR Master Mix (Promega) with a LightCycler480 (Roche). Transcript levels were calculated with the 2^−ΔCt^ method and normalized to the constitutively expressed UBQ genes. The primers are listed in Supplementary Table 1.

### Recombinant protein expression and purification

To express recombinant 6×His-SUMO-PCK1 protein, the coding sequence of PCK1 was amplified from Arabidopsis cDNA and cloned into a RSFDuet-SUMO vector with a 6xHis-SUMO tag at the N-terminus^69^ using Gibson Assembly Master Mix (E2611, NEB). To make the plasmids for expressing 6×His-SUMO-PCK1^AA^ and 6×His-SUMO-PCK1^DE^, the RSFDuet-SUMO vector containing *AtPCK1* coding sequence was mutated via Q5 Site-Directed Mutagenesis Kit (NEB, E0552). The plasmids carrying the coding sequence of *PCK1*, *PCK1^AA^* or *PCK1^DE^* were transformed into Rosetta *DE3.1* strain for protein expression. The transformed bacteria cells were cultured to OD 600 between 0.8 and 1.0 at 37°C, cooled to 16 °C, followed by addition of 300 μM isopropyl-β-D-thiogalactoside (IPTG) to induce protein expression overnight. Cell pellets were collected and resuspended in 1×Phosphate-buffered saline (PBS) buffer (2.7 mM KCl, 2 mM KH_2_PO4, 137 mM NaCl, 10 mM Na_2_HPO_4_, and pH = 7.4). The proteins were then affinity purified with Ni-NTA agarose (Invitrogen, R901-01). The expression and purification of recombinant Glutathione S-transferase (GST) tagged GST-BIN2, GST-BIN2^K69R^ and GST was performed using Glutathione agarose (Pierce, 16100) as described previously^6,70^. The affinity purified proteins were further purified by gel filtration chromatography with a Superdex 200 column (GE Healthcare). All the purified proteins were stored in 1×PBS buffer, aliquoted and kept at -80°C freezer before use. To remove the 6xHis-SUMO tag, the 6xHis-SUMO-PCK1 recombinant proteins were incubated with the SUMO protease Ulp1 (2 μg protein per unit Ulp1, Thermo Fisher Scientific, 12588018) at room temperature for 2 hours. Ulp1 and the 6xHis-SUMO fragments were then removed by incubating with the Ni-NTA magnetic beads (Thermo Fisher Scientific, 88832) at room temperature for 1 hour.

### Immunoblotting

Tissue powder (100 mg) was homogenized in 200 µl of Laemmli buffer [125 mM Tris-HCl pH 6.8, 10% (v/v) 2-mercaptoethanol, 0.01% (w/v) bromophenol blue, 4% (w/v) SDS, and 20% (v/v) glycerol] and incubated at 95°C for 10 min. After centrifugation, supernatants were separated by SDS-PAGE and blotted onto a polyvinylidene difluoride (PVDF) membrane by half-dry transfer. The PVDF membrane was blocked with 5% (w/v) low-fat milk powder (AppliChem) in TBS-T (0.1% Tween-20) at room temperature for 1 h, then probed with anti-PCK1^[52]^ (1:5,000), anti-GFP (1:2,000; HT801, Transgen Biotech), anti-His (1:3,000; 27E8, Cell Signaling Technology), or anti-GST (1:3,000; 26H1, Cell Signaling Technology), followed by washing with TBS-T for 10 min × 3 times. Subsequently, the membrane was incubated in the secondary antibody: goat anti-rabbit IgG/HRP (1:5,000, Fisher Scientific) or goat anti-mouse IgG/HRP (1:5,000, Fisher Scientific), followed by washing with TBS-T for 10 min × 3 times. Signals were detected with a WesternBright Quantum kit (Advansta).

To detect phosphorylated PCK1, proteins were separated on a 7.5% SDS-PAGE gel containing 5 mM Phos-tag™ Acrylamide (AAL-107, FUJIFILM) and 10 mM MnCl_2_. The gel was soaked in transfer buffer (Pierce, 84731) containing 1 mM ethylene diamine tetraacetic acid (EDTA) for 10 min and again soaked in transfer buffer without EDTA with gentle agitation for 10 min. Then the gel was subjected to PVDF membrane transfer and immunoblotting with antibody as described above.

### *In vitro* phosphorylation assay

For the *in vitro* phosphorylation assay, 1.0 μg recombinant GST-BIN2 protein and 1.0 μg recombinant 6×His-SUMO-PCK1 or PCK1 protein were incubated in kinase reaction buffer (25 mM Tris at pH 7.4, 12 mM MgCl_2_, 1 mM DTT, and 1 mM ATP) at 25°C for 3 h.

### Transient co-expression assay in *N. benthamiana*

*Agrobacterium tumefaciens GV3101* cells transformed with plasmids were resuspended with the induction buffer (10 mM MES pH 5.6, 10 mM MgCl_2_, 0.01% TWEEN20, and 150 μM Acetosyringone) to an absorbance OD600 of 1.0, mixed according to the combination of plasmids, and incubated for 1 h at room temperature. *Agrobacterium* cells were resuspended and infiltrated into the abaxial leaves of *N. benthamiana* using a 1 ml syringe. Two to three days after infiltration, the leaves were used analysis of the expressed proteins.

### Co-immunoprecipitation assay

The *35S:GFP/Col-0* and *pBIN2: BIN2-GFP/Col-0* stable transgenic plants were used for testing the interaction between PCK1 and BIN2 with co-immunoprecipitation (Co-IP) assay as described before^24^. Briefly, 200 mg of leaves from 4-week-old transgenic plants were harvested and grounded to fine powder in liquid nitrogen. The total proteins were extracted by the lysis buffer [50 mM Tris-HCl at pH 7.5, 150 mM NaCl, 5 mM EDTA at pH 8.0, 0.1% Triton X-100, 0.2% NP-4 with freshly added 2 mM PMSF (phenylmethylsulphonyl fluoride) and 1× protease inhibitor cocktail (Roche, 11873580001)]. After centrifugating at 4°C for 10 min, Extracts were incubated with anti-GFP magnetic beads (SM038005, Smart Lifesciences) for 1.5 h at 4°C. The precipitated samples were washed three times using the lysis buffer, eluted by boiling in Laemmli buffer for 10 min, and analyzed by anti-GFP and anti-PCK1 immunoblots as described previously.

### BLI assay

The Bio-layer interferometry (BLI) experiment was performed using a Gator-plus instrument (Gator Bio) as described previously^71^. Briefly, 20 μg/mL of purified recombinant 6xHis-SUMO-PCK1 protein was immobilized onto anti-His biosensors (Gator bio #160009). The anti-His sensors were then dipped into wells containing different concentrations of purified recombinant GST-BIN2 or GST protein as a control. Upon completion of association for 300 seconds, the sensors were dipped into 1xPBS for dissociation. The dissociation constant *K*d was generated using Gator Bio data analysis software based on the classical kinetics method (*K*_off_/*K*_on_ ratio).

### PCK1 enzymatic activity assay

Measurement of PCK1 activity in crude extracts of plant tissues was performed as described, with some modifications^61^. In brief, the harvested plant tissues were ground into fine powder with liquid nitrogen. Twenty mg grinded tissues were extracted by 500 μl Bicine-KOH (100 mM, pH 9.0) containing 10% (v/v) glycerol, 0.1% (v/v) Triton X-100, 5 mM 2-mercaptoethanol, 1 mM EDTA, 1 mM EGTA, 2 mM phenylmethylsulfonyl fluoride (PMSF), 1× protease cocktail (Roche, 11873580001), and 1× phosphatase inhibitor (Pierce, A32957). After adding the extraction buffer, samples were vortexed and then centrifuged at 14, 000 g for 10 min. Decarboxylase activity of the supernatant was measured spectrophotometrically at 340 nm in buffer [65 mM Tris-acetate (pH 7.4), 100 mM KCl, 0.3 mM OAA, 1 mM ATP, 10 μM MnCl_2_, 4mM MgCl_2_, 71.5 mM 2-mercaptoethanol, 0.4 mM NADH, 1 units of pyruvate kinase, and 2 units of lactate dehydrogenase] in a continuous assay at 25°C^[46,72]^. One unit of PCK1 activity corresponds to the production of 1 μmol product min^−1^ at 25°C. Measurement of PCK1 activity of recombinant 6×His-SUMO-PCK1, 6×His-SUMO-PCK1^AA^, and 6×His-SUMO-PCK1^DE^ was performed using 0.5 μg of recombinant proteins. The *in vitro* phosphorylated PCK1 proteins were cleaned up using Zeba Spin Desalting column (Thermo Fisher Scientific) before enzymatic assays. To measure the activity of PCK1-GFP expressed transiently in *N. benthamiana* leaves, the proteins were immunoprecipitated using a home-made anti-GFP-6xHis nanobody and HisPur Ni-NTA Magnetic Beads (Thermo Fisher Scientific) and eluted with PBS buffer containing 250 mM imidazole (Sigma-Aldrich).

### Size exclusion chromatography

Size exclusion chromatography was performed on an ÄKTA pure™ chromatography system (Cytiva) equipped with a size exclusion column, Superdex^®^ 200 Increase 10/300 GL (Cytiva, 28990944). The column’s fractionation range is from 10 kDa to 600 kDa. Samples were loaded through a 500 μL capillary loop. During the procedure, the buffer (1xPBS) flow rate was set to 0.25 mL/min, and the column was equilibrated with 2 x column volume (CV) of buffer before the sample was injected. A calibration curve was generated by plotting *K*av values versus log (molecular mass) of protein standards (CellMosaic, CM92004), including thyroglobulin (669 kDa), ferritin (440 kDa), aldolase (158 kDa), ovalbumin (43 kDa), carbonic anhydrase (29 kDa) and RNase A (13.7 kDa). *K*av values were calculated as (*V*e - *V*_0_) / (*V*_t_ - *V*_0_), where *V*e is the elution volume of the protein, *V*_0_ is the elution volume of Blue Dextran (Sigma-Aldrich, D4772) and *V*_t_ is the total volume of the column.

### Native PAGE

Native PAGE electrophoresis was performed by using the NativePAGE Bis-Tris Mini Protein Gels, 3-12% (Thermo Scientific, BN1001BOX) following the manufacturer’s instructions. A NativeMark™ unstained protein standard (Thermo Scientific, LC0725) was used as a calibration reference to estimate molecular masses of the protein bands separated by the native PAGE gels. The migration distance of each marker band and each sample band was measured as the distance from the top of the resolving gel to the center of the band using ImageJ. A standard curve was generated by linear regression of log(molecular mass, kDa) versus migration distance for the NativeMark bands (1,236, 1,048, 720, 480, 242, and 146 kDa). Molecular masses of sample bands were then determined by interpolation from the calibration curve.

### Mass spectrometry analysis

To identify the BIN2 phosphorylation sites of PCK1 *in vitro*, proteins in kinase reactions were resolved by SDS–PAGE followed by CBB staining. The gel slices containing protein bands corresponding to 6×His-SUMO-PCK1 were excised, reduced, alkylated, and digested by trypsin for LC-MS/MS analyses.

To identify *in vivo* phosphorylation sites of PCK1, two grams of 16-d-o transgenic *35S:PCK1-YFP/pck1* plants were harvested and ground to fine powder in liquid nitrogen. The total proteins were extracted by the lysis buffer (50 mM Tris-HCl at pH 7.5, 150 mM NaCl, 5 mM EDTA at pH 8.0, 0.1% Triton X-100, 0.2% NP-40) with freshly added 2 mM PMSF (phenylmethylsulphonyl fluoride) and 1 × protease inhibitor cocktail (Roche, 11873580001). After centrifugating at 4°C for 10 min, the extracts were incubated with anti-GFP magnetic beads (SM038005, Smart Lifesciences) for 1.5 h at 4°C. The beads were washed three times using the lysis buffer and eluted in 2X SDS sample buffer by boiling for 10 min. Samples were resolved by SDS–PAGE followed by Coomassie Brilliant Blue (CBB) staining. The gel slices containing protein bands corresponding to PCK1-YFP were excised, reduced, alkylated, and digested by trypsin for LC-MS/MS analyses.

LC-MS/MS analysis was performed on an Orbitrap Eclipse Tribrid mass spectrometer (Thermo Fisher), equipped with an Easy LC 1200 UPLC liquid chromatography system (Thermo Fisher). Peptides were first trapped using a trapping column (Acclaim PepMap 100 C18 HPLC, 75 μm particle size, 2 cm bed length), then separated using analytical column AUR3-25075C18, 25CM Aurora Series Emitter Column (25 cm x 75 µm, 1.7 µm C18) (IonOpticks). The flow rate was 300 nL/min, and a 120-min gradient was used. Peptides were eluted by a gradient from 3 to 28 % solvent B (80 % acetonitrile, 0.1 % formic acid) over 106 min and from 28 to 44 % solvent B over 15 min, followed by a short wash (9 min) at 90 % solvent B. Precursor scan was from mass-to-charge ratio (m/z) 375 to 1600 (resolution 120,000; AGC 200,000, maximum injection time 50ms, Normalized AGC target 50%, RF lens(%) 30) and the most intense multiply charged precursors were selected for fragmentation (resolution 15,000, AGC 5E4, maximum injection time 22ms, isolation window 1.4 m/z, normalized AGC target 100%, include charge state=2-8, cycle time 3 s). Peptides were fragmented with higher-energy collision dissociation (HCD) with normalized collision energy (NCE) 27. Dynamic exclusion was enabled for 30s.

MS/MS data was searched using Protein Prospector (v6.4.9) against the Arabidopsis *thaliana* TAIR10 protein database concatenated with randomized versions of each protein. A precursor mass tolerance was set to 10 ppm and MS/MS2 tolerance was set to 20 ppm. Carbamidomethyl on cysteine was specified as a constant modification, while the variable modifications include protein N-terminal acetylation, peptide N-terminal Gln conversion to pyroglutamate, Met oxidation, Ser, Thr, Tyr Phosphorylations. The false discovery rate (FDR) was set at 1% for both proteins and peptides. Up to two missed cleavages and a maximum of three modifications were allowed.

### Confocal microscopy imaging

Confocal microscopy imaging was performed as described previously^73^. To examine PCK1, PCK1^AA^, and PCK1^DE^ subcellular localization, roots of 8-day-old seedlings of *PCK1-YFP/pck1*, *PCK1^AA^-YFP/pck1*, and *PCK1^DE^-YFP/pck1* were stained with 0.01 g l^−1^ propidium iodide (Sigma-Aldrich) for 1 min, then briefly washed with ultrapure water. YFP florescence was detected using a Leica SP8 confocal laser scanning microscope, with an excitation wavelength at 488 nm and detection using YFP settings for YFP fluorescence or propidium iodide settings for cell wall staining. Three T3 seedlings were imaged for each of the complementation lines, alongside Col-0 as a negative control.

### Statistical analysis

All statistical analyses were done as described before^74^ with IBM SPSS Statistics (Version 29.0.2.0). All data shown in graphs and tables are presented as mean ± s.d. or as box plots, as indicated in the figure legends. Comparisons of means were performed by two-tailed Student’s *t*-test or one-way ANOVA with Tukey’s test. Different letters or asterisks represent significant differences according to *p*-values as indicated in the figure legends.

## Data availability

All materials and data presented in this study are available from the corresponding author on request. Mass spectrometry data are available via ProteomeXchange with identifier PXD075166.

## Acknowledgements

The authors thank Alberto A. Iglesias (Instituto de Agrobiotecnología del Litoral, UNL, CONICET, FBCB, Santa Fe, Argentina) for sharing the anti-PCK1 antibody, and Andres Reyes for mass spectrometry data analysis. This work was funded by the National Institutes of General Medical Science grants R01GM066258 to Z-Y.W and S10OD030441 to S.-L.X. and by the Carnegie Endowment Fund to the Carnegie Mass Spectrometry Facility.

## Contributions

H.Z. and Z.W. designed the research. H.Z., Y.A., K.U., A.C., C.T. performed experiments. H.Z., S.X., and Z.W. analyzed data. H.Z. and Z.W. wrote the manuscript. All authors edited the manuscript.

## Competing interests

The authors declare no competing interests.

---

**Extended Data Fig. 1:**
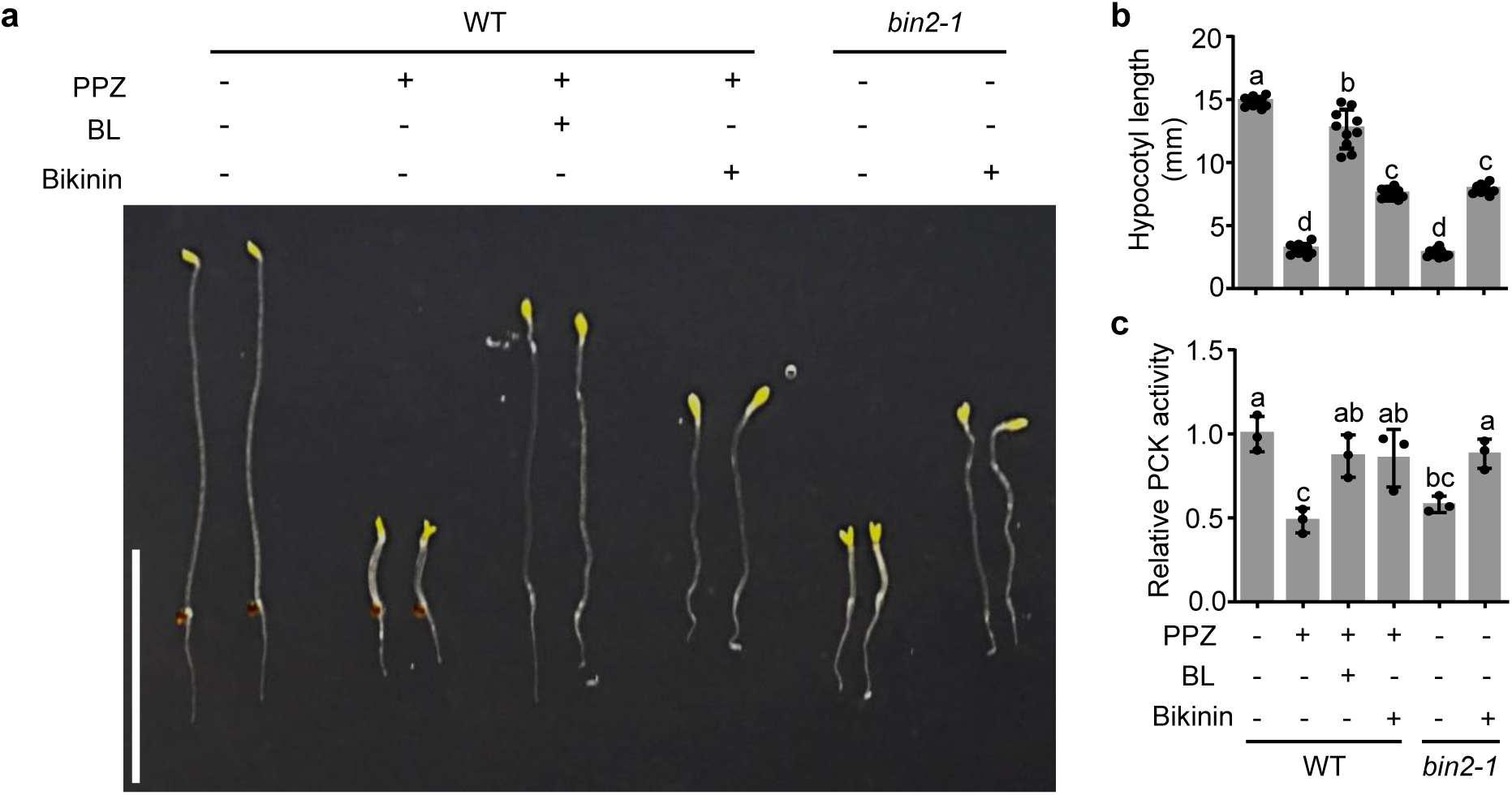
BRs increase PCK activity and hypocotyl elongation. **a,** Photograph of 5-day-old wild-type (WT) and *bin2-1* seedlings grown on 0.5x MS sugar-free media containing 2 μM propiconazole (PPZ), 100 nM brassinolide (BL), and 30 μM bikinin. Scale bar, 10 mm. **b,c,** Hypocotyl length (**b**) and relative PCK decarboxylation activity (**c**) of seedlings shown in (**a**). Data are mean ± s.d., *n* = 10 seedlings (**b**), *n* = 3 biological replicates (**c**); Significant differences between means are marked by different letters (one-way ANOVA, Tukey’s test, *P <* 0.05).

**Extended Data Fig. 2:**
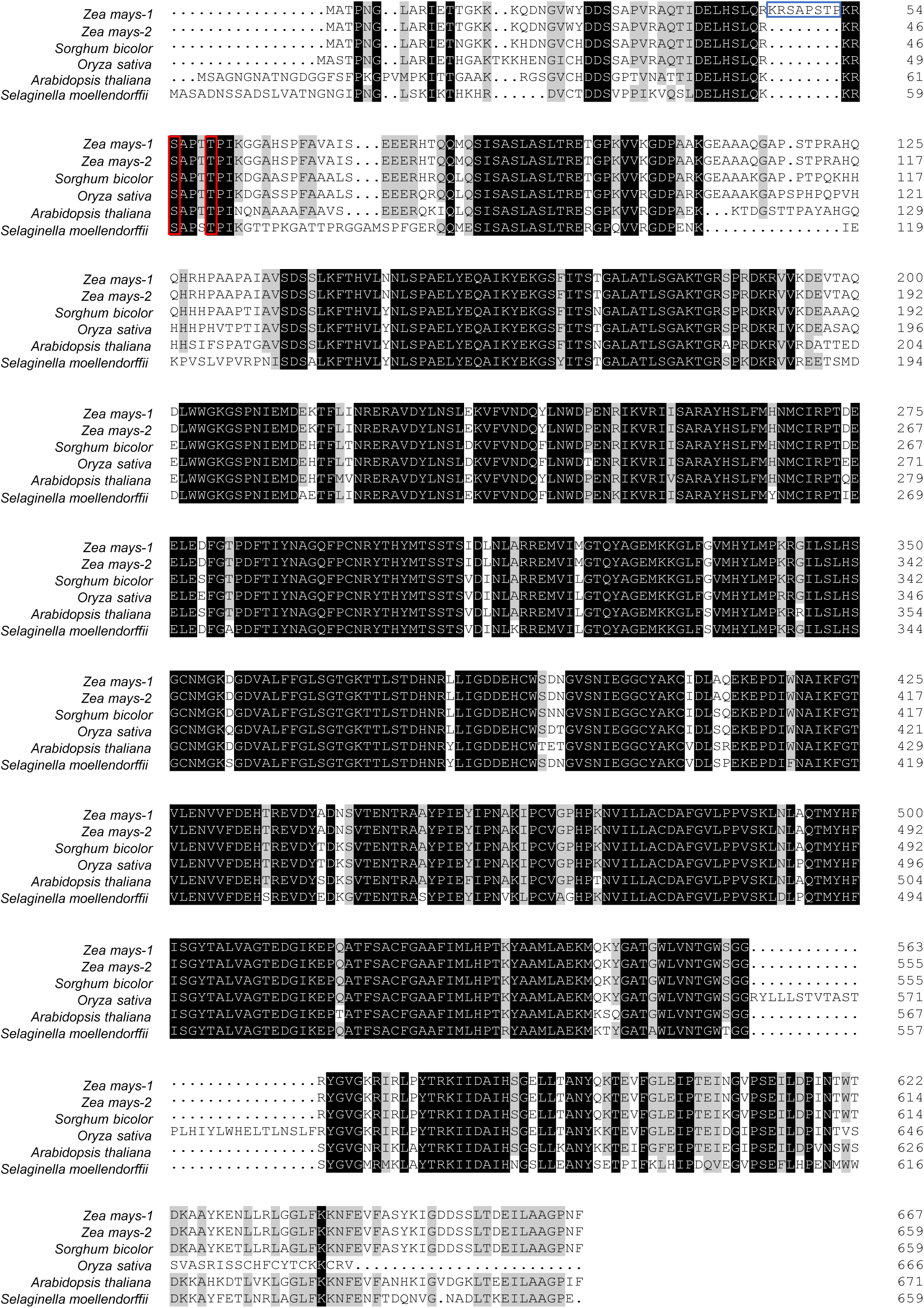
Sequence alignment of PCK homologues in plant species. The two PCK isoforms in *Zea mays* are designated *Zea mays-1* and *Zea mays-2*. Two conserved Ser/Thr residues (Ser-62 and Thr-66 of AtPCK1) are marked by red boxes. The repeat peptide sequence (45-KRSAPSTP-52) of *Zea mays-1* is marked by a blue box.

**Extended Date Fig. 3:**
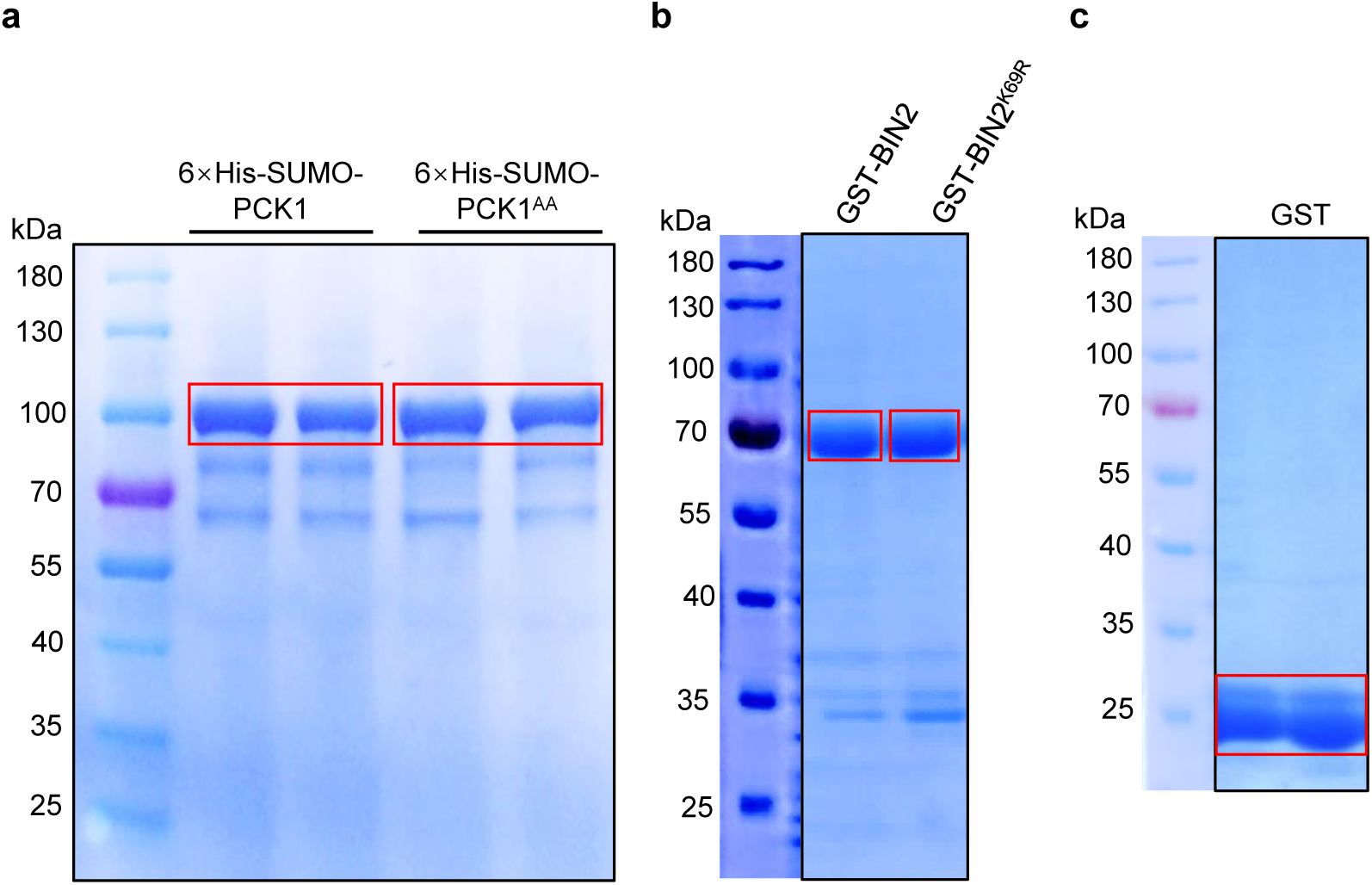
SDS-PAGE analysis of the 6×His-SUMO-PCK1 and 6×His-SUMO-PCK1^AA^ (**a**), GST-BIN2 and GST-BIN2^K69R^ (**b**), and GST (**c**) proteins purified from *E.coli*.

**Extended Data Fig. 4:**
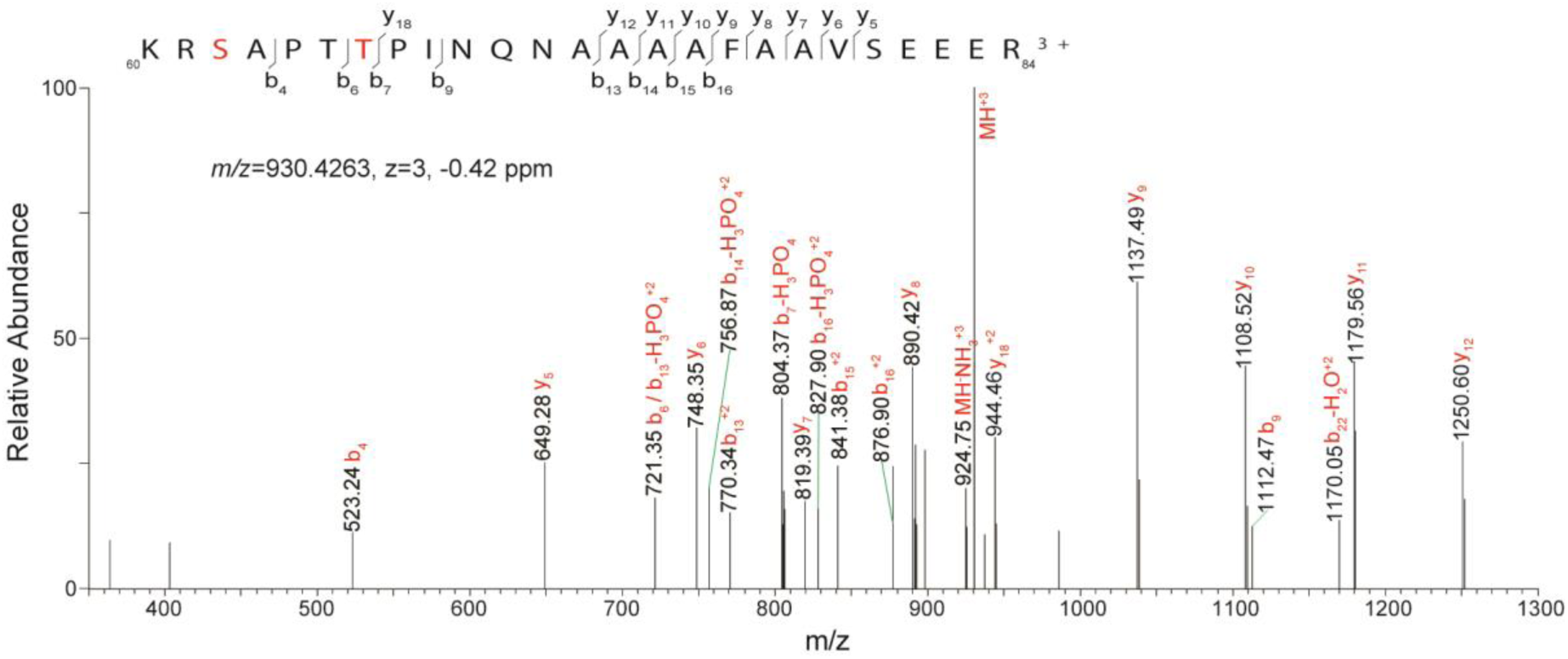
An MS/MS spectrum of phospho-peptide 60-KRSAPTTPINQNAAAAFAAVSEEER-84, identifying phosphorylation on Ser-62 and Thr-66 (highlighted in red) of recombinant His-SUMO-PCK1 from the *in vitro* kinase assay in (Fig. 2c).

**Extended Data Fig. 5:**
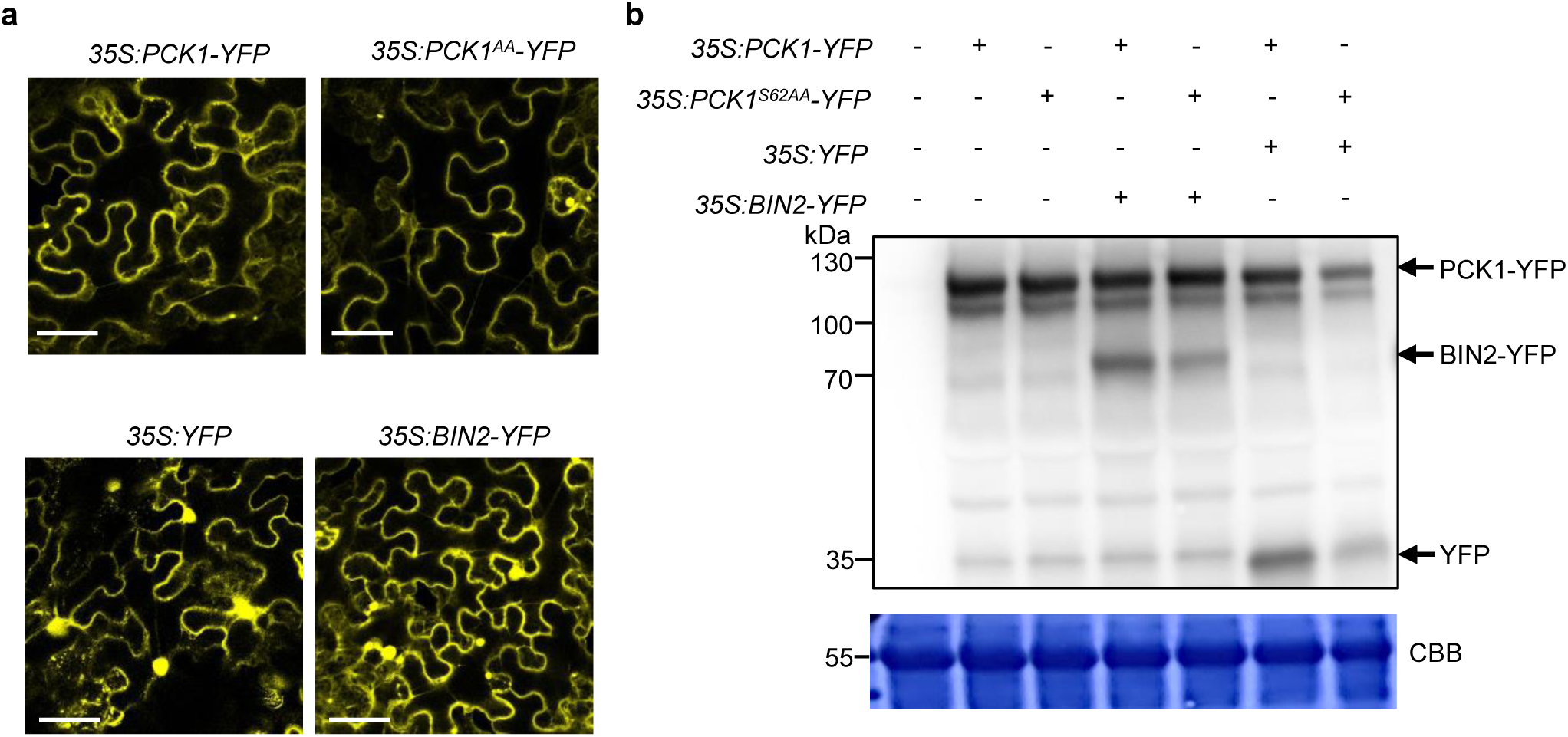
PCK1 is phosphorylated by BIN2 *in vivo*. **a,** Representative confocal laser scanning microscopic images of epidermal cells of 6-week-old *N. benthamiana* leaves infiltrated with *Agrobacterium tumefaciens* carrying the indicated plasmids (related to Fig. 2d). Scale bars, 50 μm. **b,** Anti-GFP immunoblot shows the expression of proteins from the infiltrated *N. benthamiana* leaves used in (Fig. 2d). CBB: Coomassie Brilliant Blue staining of the gel blot.

**Extended Data Fig. 6:**
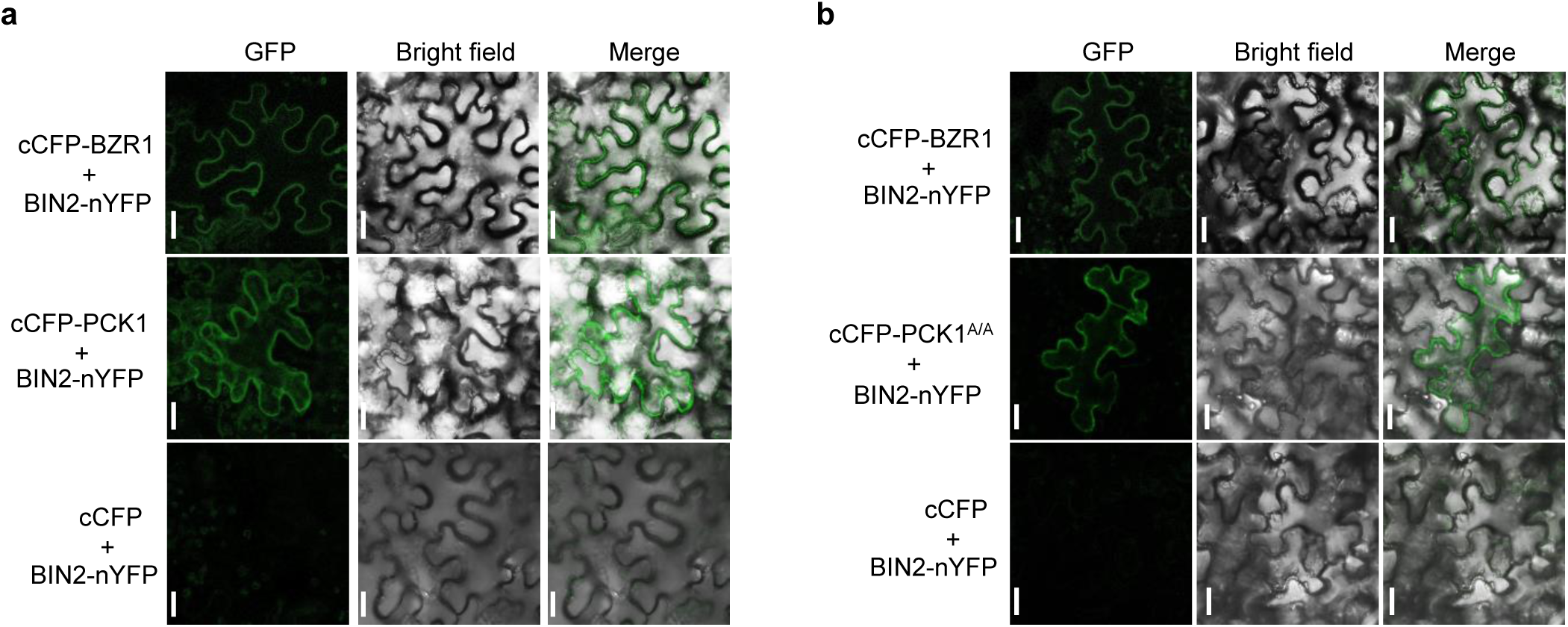
Bimolecular fluorescence complementation (BiFC) analyses of BIN2 interaction with PCK1. (**a**) **and PCK1^AA^** (**b**). Representative confocal images of *N. benthamiana* leaf epidermal cells transiently transfected with the indicated constructs. The known interaction between BZR1 and BIN2 was used as a positive control. Scale bars, 20 μm.

**Extended Data Fig. 7:**
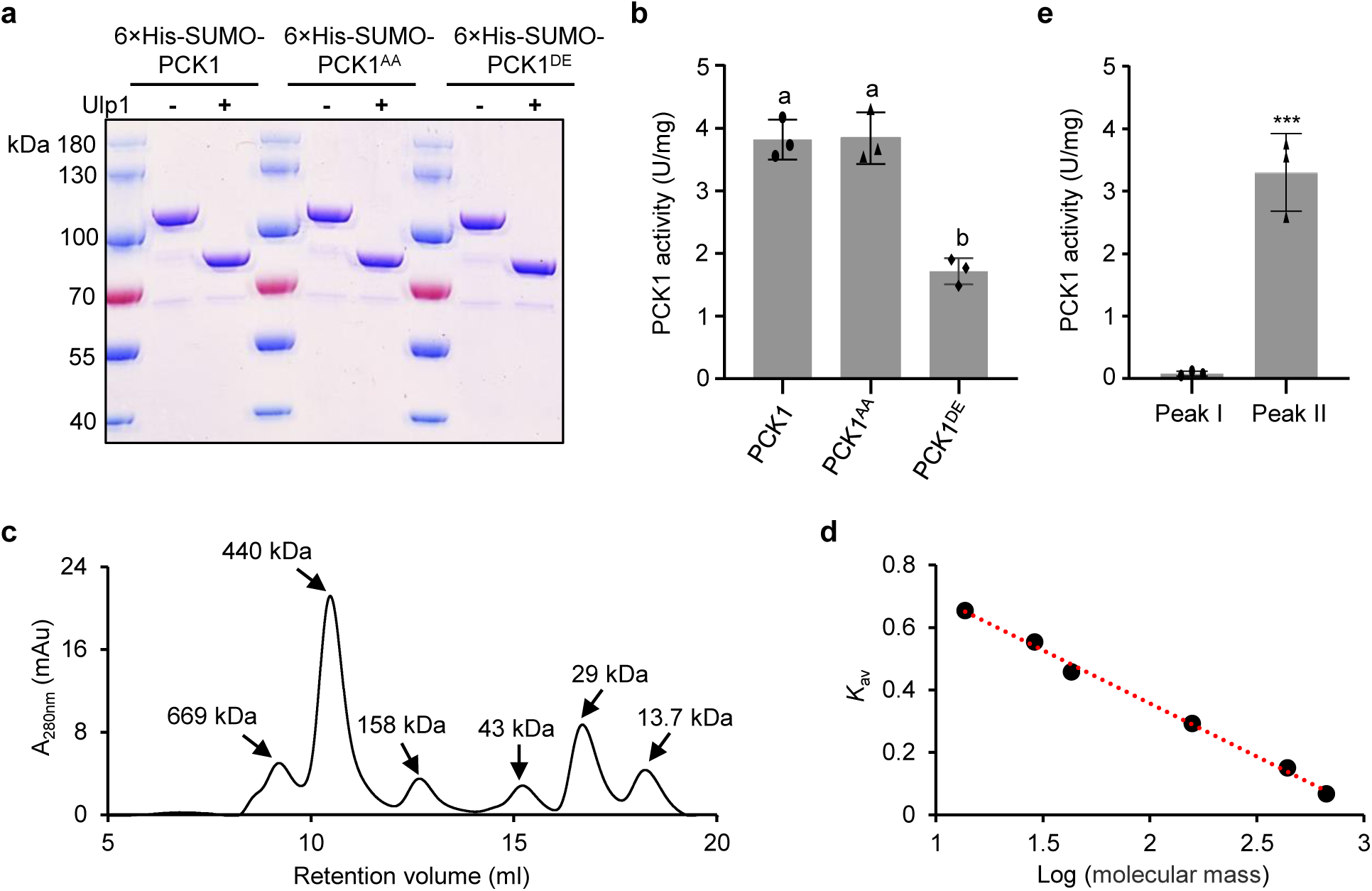
Characterization of recombinant PCK1 proteins. **a,** SDS-PAGE analysis of recombinant proteins before and after cleavage by the sumo protease Ulp1. **b,** Activities of the recombinant PCK1, PCK1^AA^, and PCK1^DE^ proteins without the 6xHis-SUMO tag. PCK activities are mean ± s.d., *n =* 3 replicates; Letters above the bars reflect significant differences between means (one-way ANOVA, Tukey’s test, *P <* 0.05). **c,** SEC spectrum of protein standard. **d,** Standard curve used to calculate the molecular mass (MM). The MM was negatively correlated with the retention volume (*y = -0.3403x + 1.0377, R²* = 0.9964). **e.** PCK decarboxylase activities of the two SEC peak fractions of PCK1 after cleavage of 6xHis-SUMO tag and phosphorylation by GST-BIN2. Data are mean ± s.d., *n* = 3 replicates; ***, *P <* 0.001, according to two-tailed Student’s *t*-test.

**Extended Data Fig. 8:**
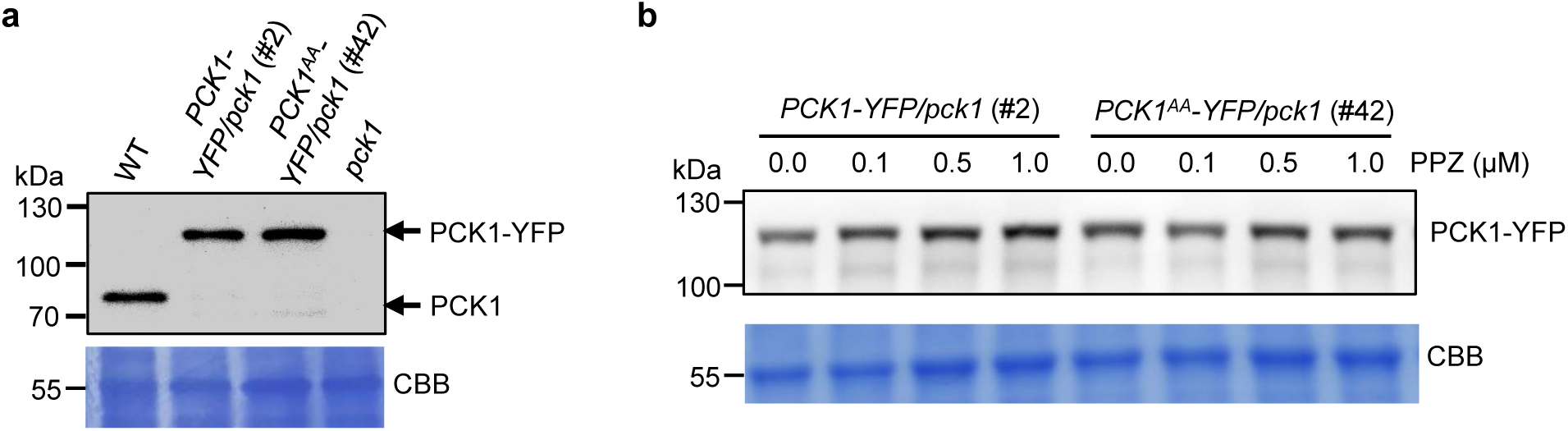
*PCK1-YFP/pck1* and *PCK1^AA^-YFP/pck1* lines expressed wild-type levels of PCK1, which were unaffected by PPZ treatment. **a,** Anti-PCK1 immunoblotting analysis of PCK1 protein levels in 5-day-old WT, *pck1* mutant, *PCK1-YFP/pck1* and *PCK1*^AA^*-YFP/pck1* seedlings grown on 0.5x MS medium in the dark. **b,** Anti-GFP immunoblotting analysis of *PCK1-YFP/pck1* and *PCK1^AA^-YFP/pck1* seedlings grown on 0.5x MS medium containing the indicated concentrations of propiconazole (PPZ). CBB: Coomassie Brilliant Blue staining of the gel blot.

**Extended Data Fig. 9:**
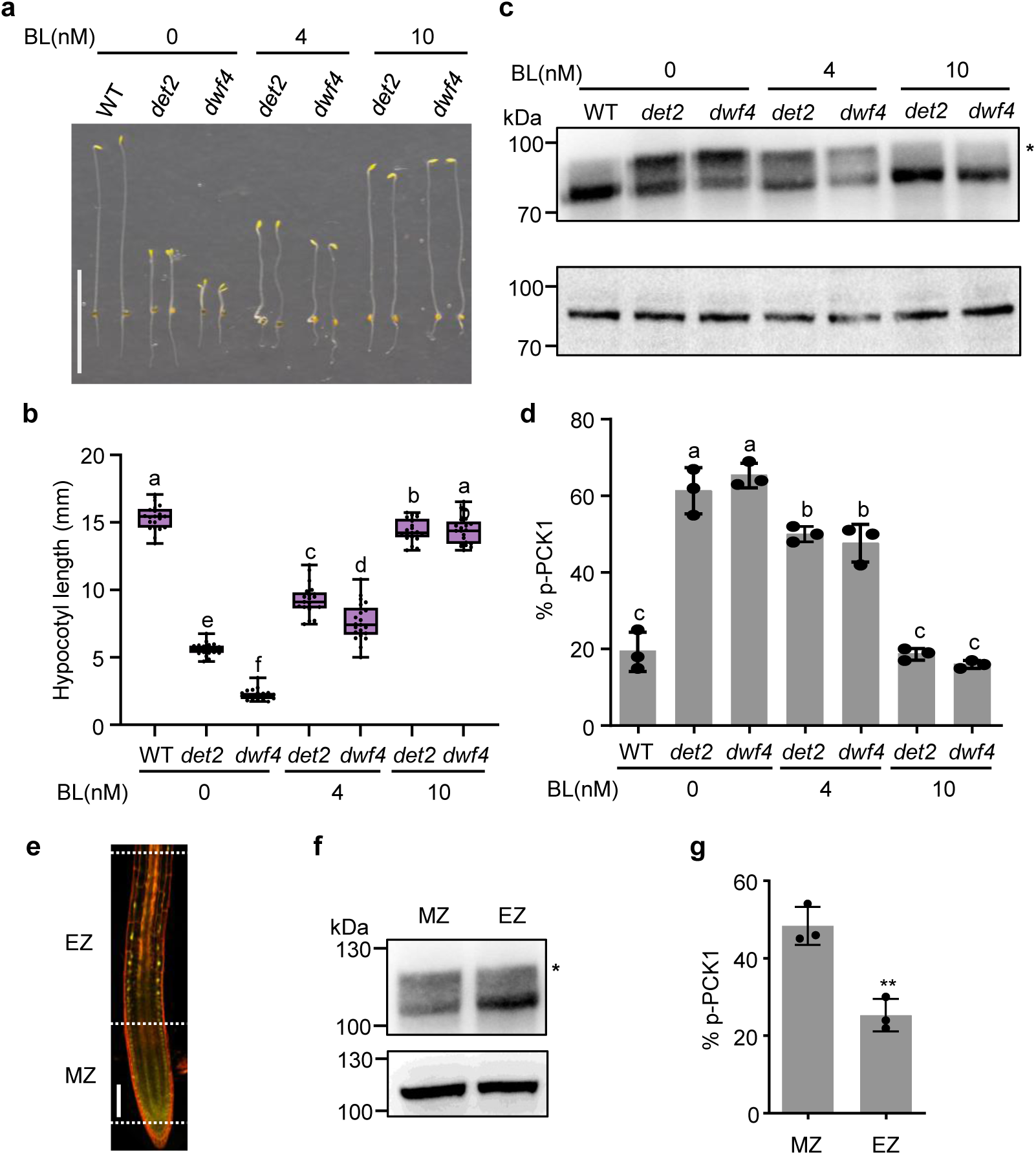
BRs regulate phosphorylation of PCK1. **a,b**, Wild-type (WT), *det2,* and *dwf4* seedlings were grown on 0.5x MS sugar-free medium containing the indicated concentrations of brassinolide (BL) in the dark for 4 days. Representative seedlings were imaged (**a**), and hypocotyl lengths (**b**) were measured. Scale bar, 10 mm. *n =* 21 seedlings, the central lines within the box plots represent the medians, the box represents the interquartile range (IQR), the whiskers extend to minima and maxima. **c,** Phosphorylation status of PCK1 of WT, *det2*, and *dwf4* seedlings shown in (**a**). Phos-tag gel blot (upper panel) or SDS-PAGE (lower panel) probed with an anti-PCK1 antibody. Phosphorylated PCK1 (p-PCK1) is the shifted band marked with *. **d,** The % of p-PCK1 was calculated from the band intensities of three replicates of the immunoblot shown in (**c**). Data are mean ± s.d. Different letters reflect significant differences between means (one-way ANOVA, Tukey’s test, *P <* 0.05). **e,** Representative images of BZR1-YFP fluorescence of root tip of *pBZR1:BZR1-YFP/Col-0* transgenic seedlings grown on 0.5x MS medium in the light for 5 days. Scale bar, 100 μm. MZ, meristem zone; EZ, elongation zone. **f,** Phosphorylation status of PCK1 in the MZ and EZ of the *PCK1-YFP/pck1* transgenic seedlings. PCK1 protein was analyzed by Phos-tag gel (upper panel) or SDS-PAGE (lower panel) gel blots probed with an anti-PCK1 antibody. Phosphorylated PCK1 (p-PCK1) is the shifted band marked with *. **g,** The % of p-PCK1 was calculated from the band intensities of three replicates of the immunoblot shown in (**f**). Data are mean ± s.d.**, *P <* 0.01, according to two-tailed Student’s *t*-test.

**Extended Data Fig. 10:**
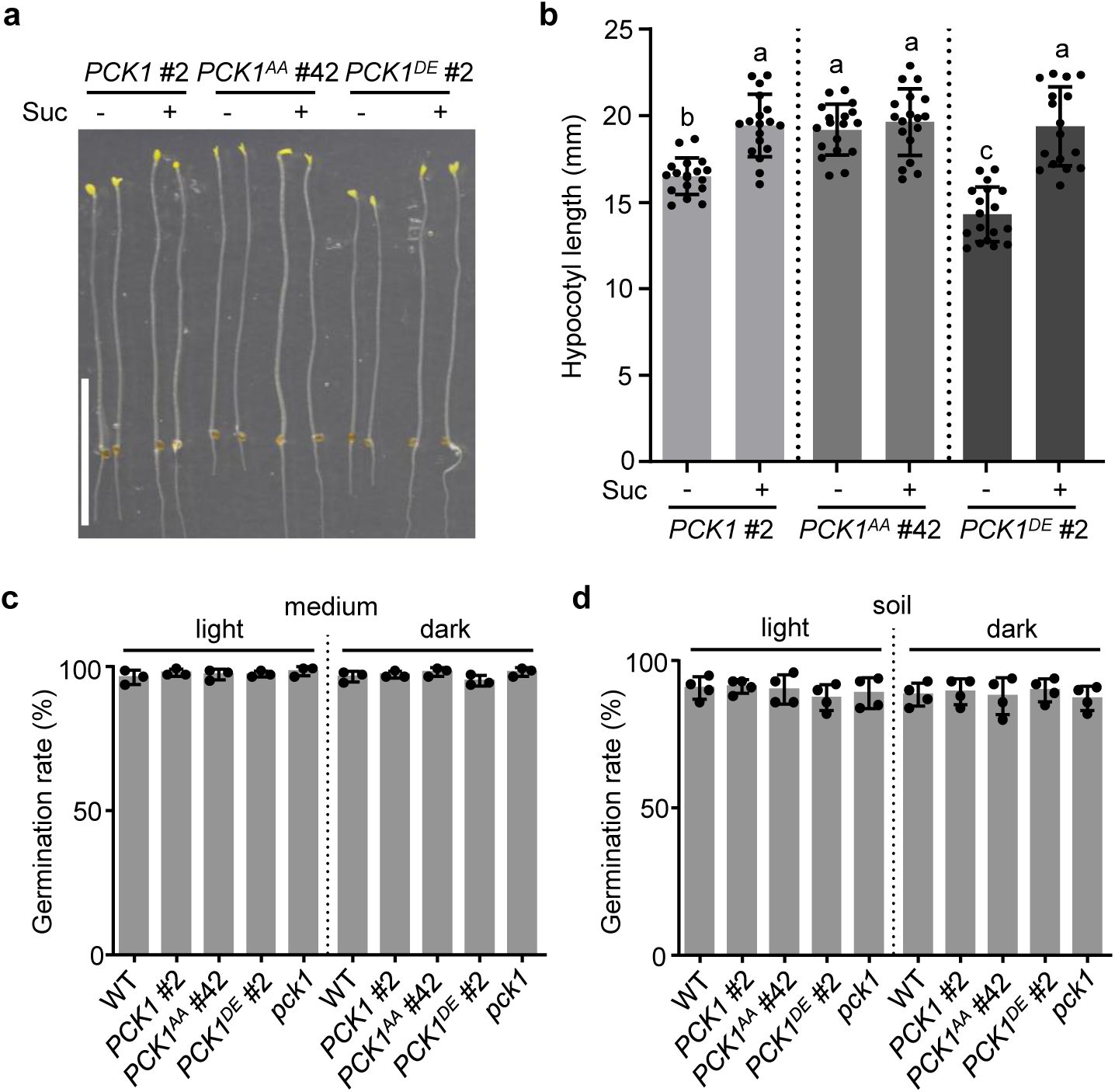
Germination and growth phenotypes of *PCK1-YFP/pck1*, *PCK1^AA^-YFP/pck1*, and *PCK1^DE^-YFP/pck1* transgenic seedlings. Representative phenotypes (**a**) and hypocotyl lengths (**b**) of *PCK1-YFP/pck1* (*PCK1* #2), *PCK1^AA^-YFP/pck1* (*PCK1^AA^*#42), and *PCK1^DE^-YFP/pck1* (*PCK1^DE^* #2) transgenic seedlings grown for 5.5 days on 0.5x MS medium with or without 1% sucrose. Scale bar, 10 mm. Data are mean ± s.d., *n* = 18 seedlings. Different letters reflect significant differences between means (one-way ANOVA, Tukey’s test, *P <* 0.05). **c,d,** Seeds of wild-type (WT), *pck1*, *PCK1-YFP/pck1* (*PCK1*), *PCK1^AA^-YFP/pck1* (*PCK1^AA^*), and *PCK1^DE^-YFP/pck1* (*PCK1^DE^*) transgenic lines were sown on 0.5x MS sugar-free medium (**c**) or on soil (**d**) in the light and dark. Germination rates of seeds were calculated after 3 days.

## Notes

### Competing Interest Statement

The authors have declared no competing interest.

### Summary of Updates

We added data to Figure 7

