## Supplemental Table 1 for "Brassinosteroids promote sugar synthesis by inhibiting BIN2 phosphorylation of phosphoenolpyruvate carboxykinase"

**Supplementary Table 1. Primers used in this study**

| Primer name | Sequence (5' to 3') |
|---|---|
| p101_PCK1-CDS-F | CACCATGTCGGCCGGTAACGGAAATGCTAC |
| p101_PCK1-CDS-R | AAAGATAGGACCAGCAGCGAGAATCTCC |
| pSUMO_PCK1-CDS-[illegible] | AGAACAGATTGGTGGATCCGGAATGTCGGCCGGTAACGGAAAT |
| pSUMO_PCK1-CDS-[illegible] | TAAGCATTATGCGGCCGCACTAAAAGATAGGACCAGCAGCGAGAATCT |
| qPCK1-F | AGCTTTGGTACTCCGGATTTTA |
| qPCK1-R | CCTAGCCAGATTAAGGTCTACG |
| qUBQ10-F | ATCACCCTTGAAGTGGA |
| qUBQ10-R | GAAACCACCACGAAGAC |
| $PCK1^{AA}$-F1 | AAACGTGCTGCTCCTACCACACC |
| $PCK1^{AA}$-R1 | GTAGGAGCAGCACGTTTCTTCTG |
| $PCK1^{AA}$-F2 | CTCCTACCGCACCGATCAAC |
| $PCK1^{AA}$-R2 | GATCGGTGCGGTAGGAGCAGAAC |
| $PCK1^{DE}$-F1 | CACCATGTCGGCCGGTAACGGAAATGCTAC |
| $PCK1^{DE}$-R1 | GTTGATCGGTTCGGTAGGAGCGTCACGTTTCTTCTGTAACGAAT |
| $PCK1^{DE}$-F2 | GCTCCTACCGAACCGATCAACCAAAACGCCGCCGCT |
| $PCK1^{DE}$-R2 | AAAGATAGGACCAGCAGCGAGAATCTCC |

| Purpose |
|---|
| Amplify PCK1 coding region to make 35S:PCK1-YFP |
| Amplify PCK1 coding region to make 6×His-SUMO-PCK1 |
| RT-qPCR of *PCK1* |
| RT-qPCR of *UBQ10* |
| To make $PCK1^{AA}$ |
| To make $PCK1^{DE}$ |
